# Single-Cell Transcriptomics Unveils Gene Regulatory Network Plasticity

**DOI:** 10.1101/446104

**Authors:** Giovanni Iacono, Ramon Massoni-Badosa, Holger Heyn

## Abstract

**SUMMARY:** Single-cell RNA sequencing (scRNA-seq) plays a pivotal role in our understanding of cellular heterogeneity. Current analytical workflows are driven by categorizing principles that consider cells as individual entities and classify them into complex taxonomies. We have devised a conceptually different computational framework based on a holistic view, where single-cell datasets are used to infer global, large-scale regulatory networks. We developed correlation metrics that are specifically tailored to single-cell data, and then generated, validated and interpreted single-cell-derived regulatory networks from organs and perturbed systems, such as diabetes and Alzheimer’s disease. Using advanced tools from graph theory, we computed an unbiased quantification of a gene’s biological relevance, and accurately pinpointed key players in organ function and drivers of diseases. Our approach detected multiple latent regulatory changes that are invisible to single-cell workflows based on clustering or differential expression analysis. In summary, we have established the feasibility and value of regulatory network analysis using scRNA-seq datasets, which significantly broadens the biological insights that can be obtained with this leading technology.

## INTRODUCTION

scRNA-seq is the leading technology for exploring tissue heterogeneity, unravelling the dynamics of differentiation, and quantifying transcriptional stochasticity. scRNA-seq data are being used to answer increasingly demanding biological questions, which has driven the development in recent years of an array of computational tools for scRNA-seq analysis (Zappia et al., 2018). Currently, these tools focus on improving features such as clustering, retrieving marker genes, and exploring differentiation trajectories (Zappia et al., 2018). These scenarios are inspired by a dividing, fragmenting principle, where each cell is an independent identity that must can be categorized into different types or stages of increasing hierarchical complexity. This is illustrated by recent large-scale cell atlases that often reach hundreds of stratified (sub)clusters (Zeisel et al., 2018). This has undoubtedly improved our understanding of cell diversity in various biological contexts. However, we hypothesize that a very different approach, inspired by a unifying rather than dividing ideal, would add a novel layer of information that would significantly increase the knowledge gained from single-cell datasets.

Gene expression is tightly regulated by networks of transcription factors, cofactors and signaling molecules. Understanding these networks is a major goal in modern computational biology, as it will allow us to pinpoint crucial factors that determine phenotype in healthy systems as well as in disease (Emmert-Streib et al., 2014; Thompson et al., 2015). Unraveling the determinants of a given phenotype provides mechanistic insights into causal dependencies in complex cellular systems. Potentially, single-cell information offers the opportunity to derive a global regulatory network (Fiers et al., 2018). Traditional approaches to transcriptome profiling, namely microarray and RNA-seq of pooled cells, have been successfully used to infer and characterize regulatory networks, with a recent example using 9,435 bulk RNA-seq samples to decode tissue-specific regulatory networks (Sonawane et al., 2017). To date, there are only small-scale efforts to derive regulatory networks from single-cell transcriptomics data, and these efforts have been restricted to specific network properties (Guo et al., 2015; Lim et al., 2016). This seems unexpected given that single-cell sequencing is the ideal technology for monitoring real interactions between genes in individual cells. However, single-cell data is undermined by a series of technical limitations, such as dropout events (expressed genes undetected by scRNA-seq) and a high level of noise, which have made it difficult to infer regulatory networks using this type of data (Chen and Mar, 2018).

In this paper, we demonstrate the feasibility and value of regulatory network analysis using scRNA-seq datasets. We present a novel correlation metric that can detect gene-to-gene correlations that are otherwise hidden by technical limitations. We apply this new metric to generate global, large scale regulatory networks for 11 mouse organs (Tabula Muris Consortium et al., 2018), for pancreas tissue from healthy individuals and patients with type 2 diabetes (Segerstolpe et al., 2016), and for a mouse model of Alzheimer’s disease (Keren-Shaul et al., 2017). We then validate the resulting networks at multiple levels to confirm the reliability of the reconstruction. Next, we analyze the networks using state-of-the-art tools borrowed from graph theory, such as node centralities and dynamical properties. Finally, we integrate network-driven results with standard analyses such as clustering and differential expression analysis, and show that key regulators of healthy and diseased systems can only be identified by using holistic, network-based approaches. Together, our results represent the first complete, validated, high-throughput and disease-centered application of single-cell regulatory network analysis, significantly increasing the knowledge gained from this leading technology.

## RESULTS

### Inferring regulatory networks from large-scale single-cell transcriptomics

We initially set out to develop a reliable approach for inferring global regulatory networks from single-cell data (**Fig. 1**). To generate a regulatory network starting from expression data, we require a robust measure of correlation between genes. Unlike in RNA-seq from pools of cells (bulk), single-cell data is inherently noisy and highly sparse, which prevents the effective use of standard metrics such as *Pearson, Spearman* or *Cosine* correlation, or even *mutual information* (Methods). Hence, we conceived a novel correlation measure based on *bigSCale* (Iacono et al., 2018), a computational framework used to analyze single-cell data. Briefly, instead of searching for relationships using the original variables, namely (normalized) expression counts, we compute the correlations between transformed variables, in which expression counts are replaced by Z-scores. These Z-scores are derived from an unsupervised analysis based on iterative differential expression (DE) between small clusters of cells (Methods). To compute Z-scores, we exploit a probabilistic model of the noise that considers all sources of variability in singlecell data. Thereby, this approach can detect correlations that would otherwise be concealed by drop-out events and other technical artifacts. When applied to a scRNA-seq dataset of 7,697 microglia cells (Keren-Shaul et al., 2017), *bigSCale* identified 933,936 significant gene-to-gene correlations (*Pearson* >0.8), a gain of almost 40,000-fold compared to normalized UMI count data (only 24 correlations, **Fig. 2A,B**). This large increase in the number of detected correlations is supported by a radically different distribution in the Z-score compared to that for the UMIs/reads space (**Fig. 2A**). When applied to seven additional datasets generated using different scRNA-seq techniques (Fluidigm C1, 10x Genomics Chromium, MARS-seq, Smart-seq2), with different sequencing depths and from different tissue sources (Iacono et al., 2018; Tabula Muris Consortium et al., 2018; Zeisel et al., 2015), the *bigSCale* metric consistently outperformed standard approaches, suggesting that it is a valid universal correlation metric for scRNA-seq data (**Fig. 2C**).

**Figure 1.**
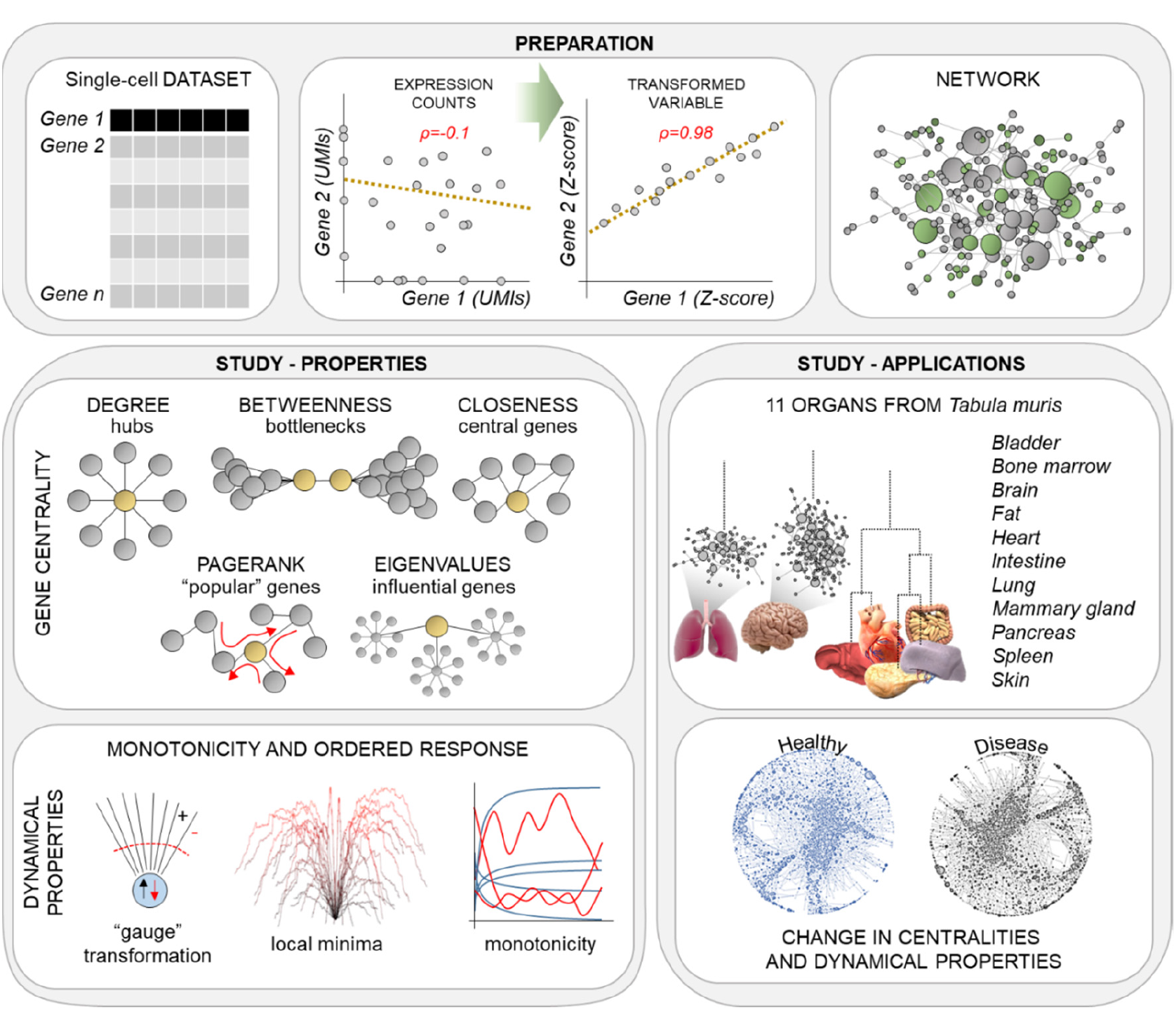
Overview of the computational framework. A change of variable (from expression values to Z-score) is used to detect otherwise hidden correlations between genes in single-cell datasets, ultimately allowing us to infer the global regulatory network. Network properties are characterized using concepts from graph theory. We generated, compared and characterized the networks of 11 organs in the mouse (*tabula muris*), of pancreas from healthy and type 2 diabetes human subjects, and of a mouse model of Alzheimer’s disease.

**Figure 2.**
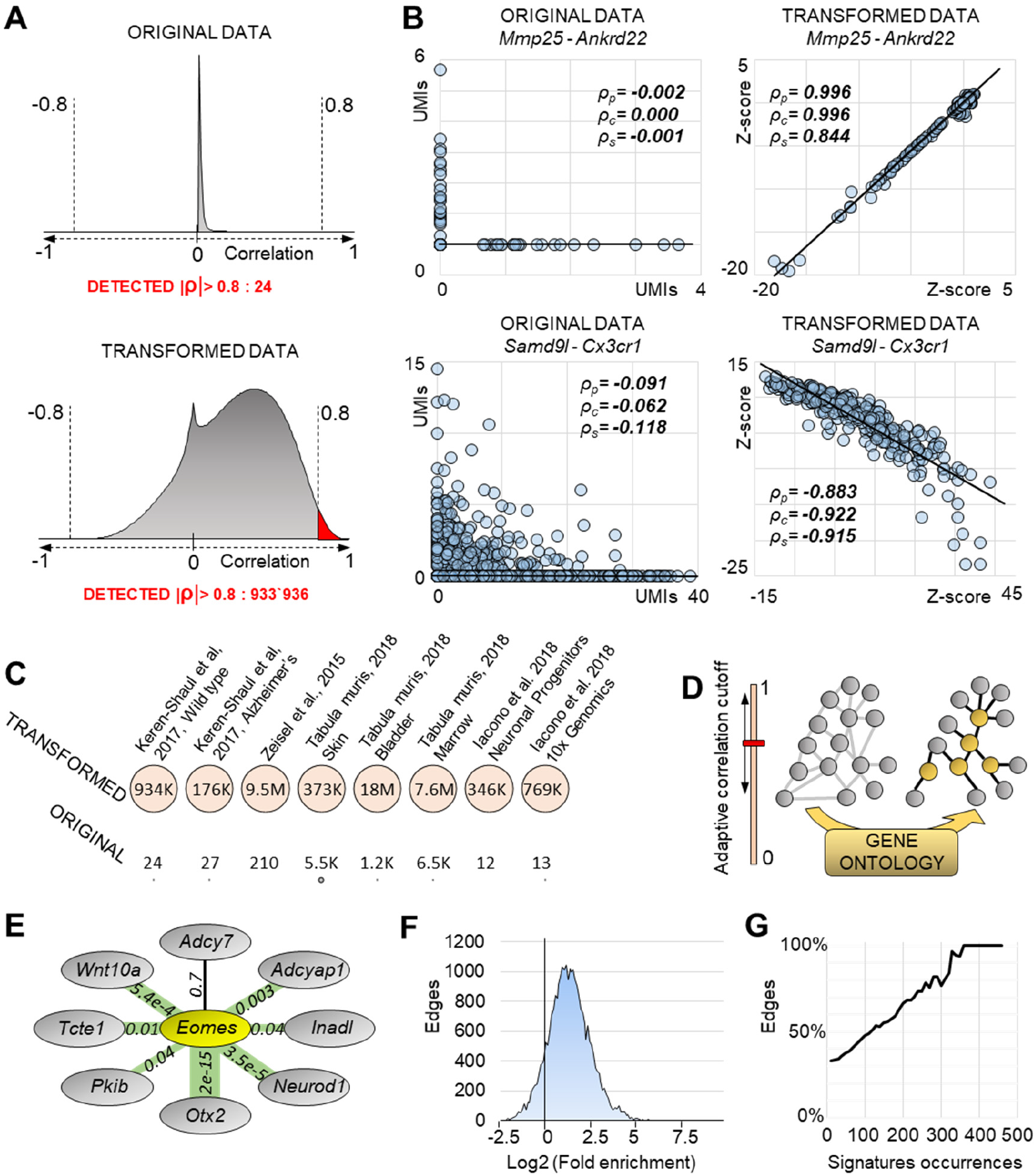
A metric tailored to single-cell data allows detection of hidden correlations. (A) Distribution of *Pearson* correlations ρ_p_ in normalized expression data (7697 microglia cells) or in the Z-score space. We detect only 24 correlations |ρ_p_|>0.8 in the first scenario, but almost one million |ρ_p_|>0.8 in the Z-score space. (B) Examples of correlations using either expression values or Z-score transformed data (ρ_p_ *Pearson*, ρc *Cosine*, ρs *Spearman*). Due to drop-out events and other artifacts, the positive correlation between *Mmp25* and *Ankrd22* is only exposed using Z-scores. Similarly for the negative correlation between *Samd9l* and *Cx3cr1.* (C) Comparison of detected correlations |ρ_p_|>0.8 using either original expression values or Z-score transformed data across different scRNA-seq technologies, sequencing depths (from 625 (Keren-Shaul et al., 2017) to 6480 (Iacono et al., 2018) average detected genes per cell) and source material. (D) An adaptive correlation cutoff and GO annotations are used to infer the regulatory networks from correlation data. (E) Example of the validation of *Eomes* neighbors in the brain network. For each edge, we compute fold enrichment (proportional to edge width, highest *Otx2* with 9.87, lowest *Adcy7* with 0.83) and a p-value (labels). In the case of *Eomes*, all edges but one (*Adcy7*) are validated with p<0.05. (F) Overall distribution of edge-wise fold enrichment in the brain network is biased towards positive values, suggesting that neighboring genes are regulated together in perturbed systems. (G) Brain network, percentage of validated edges with p<0.05 (y-axis) filtered by the number of GSEA occurrences (x-axis). Some genes have few occurrences, which are less likely to yield significant p-values in the *Fisher’s* exact test. Higher percentages of validated (p<0.05) edges are obtained by considering only edges with high occurrences.

After identifying gene-to-gene correlations, an adaptive threshold is applied to retain only significant correlations (Methods). This adaptivity equalizes the effects of different cell numbers and coverage, and other technical features of scRNA-seq datasets. The retained correlations then become the weighted edges of the regulatory network, with either positive or negative signs. In the final step, gene ontology (GO) information is used to subset the network to “*regulators of gene expression*”, in order to retain only putative causal (regulatory) relationships (Methods) (**Fig. 2D**). Note that using external information (e.g. GO) is an established method for refining networks (Cheng et al., 2009; Tuncay et al., 2007; Zhang et al., 2008). To determine the importance of a given gene in a single-cell regulatory network and its underlying biological system, we applied advanced analytical tools from the field of graph theory. These tools allow us to quantify the biological relevance of a gene using various measures of centrality, namely *degree*, *betweenness, closeness, pagerank* and *eigenvalues* (**Fig. 1**). For example, *betweenness* centrality identifies regulatory bottlenecks, i.e. genes crucial for the flow of information; *closeness* centrality identifies genes situated in a central position in a network and *eigenvalue* centrality indicates highly influential genes whose signal quickly spreads throughout the network.

### Single-cell regulatory networks identify essential and specific genes for organ function

To evaluate the value of using large-scale regulatory networks inferred from single cells to aid biological interpretation of scRNA-seq datasets, we first applied our framework to a single-cell resolved mouse organ atlas (Tabula Muris Consortium et al., 2018). We generated regulatory networks from 11 organs: endoderm (lung, pancreas, intestine), mesoderm (heart, fat, spleen, bladder, bone marrow) and ectoderm (skin, brain, mammary glands). The adaptive correlation threshold required to normalize batch effects such as sequencing depth or cell numbers reached high values for all organs (ρ_thresh_>0.9, **Table 1**), which confirms the significance of selected correlations. Inferred networks had a scale-free topology (a structure conferring fault-tolerant behavior, **Fig. S1**), which is in line with previous findings in manually curated networks (Albert, 2005; Balaji et al., 2006; Noort et al., 2004). We next sought to validate our predicted regulatory edges. We reasoned that when the system is perturbed all pairs of genes linked by an undirected edge should be activated/deactivated together. Thus, we used the Gene Set Enrichment Analysis (GSEA) database, which contains an extensive collection of experimental signatures representing perturbations in different biological systems (Methods). We performed a proportional test (*Fisher’s* exact test) to quantify the co-occurrence of neighboring genes in GSEA experimental signatures, thereby testing the significance of each individual edge in the network (**Fig. 2E**). In the brain network, the edges (23,492) showed an overall distribution bias toward positive fold-enrichment and significant p-values, which supports our inferred regulatory links (**Fig. 2F,G**). Specifically, 34% of the edges were validated (p<0.05), and this percentage increased when we considered only the edges whose genes are present in many GSEA signatures (**Fig. 2G**). In fact, 100% of the edges were validated when considering only genes appearing in at least 360 signatures. The results were similar for the other 10 mouse organ networks (**Fig. S2A**).

**Table 1.**
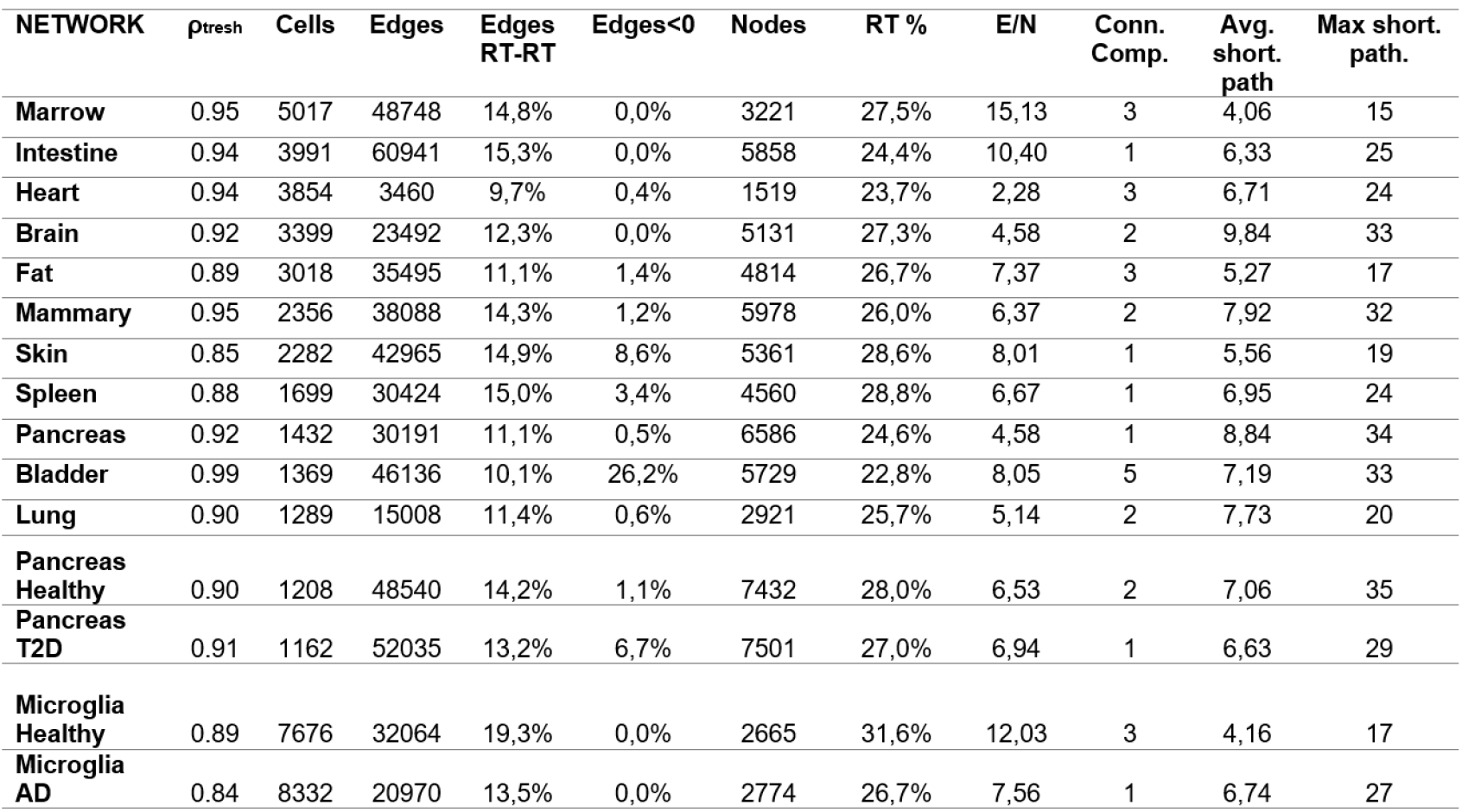
Overview of specifications for inferred regulatory networks. In order: the adaptive correlation threshold set to retain significant correlations. Number of cells, edges. Percentage of edges between regulators of transcription (RT) and of edges with negative sign. Amount of nodes and their percentage being RT. Ratio edges/node (E/N), number of connected components, average and maximum shortest paths.

We found that specific global properties differed markedly across organ-specific networks while other properties remained largely homogeneous (**Table 1**). For example, we observed diverse signaling complexities (ratio edges/node), this being lowest in the heart (2.28 edges per node) and highest in bone marrow (15.13 edges per node). In contrast, the percentage of “regulators of gene expression” (nodes actively regulating the expression of other nodes) was very stable across all tissues (26-28%). Notably, genes that were central in the different measures showed marginal overlap (**Fig. 3A, Fig. S3,4**), which suggests that conceptually different centralities quantifies distinct types of biological importance and provide mutually complementary information. To confirm the importance of central regulatory genes in the biological system, we calculated their enrichment among experimentally validated essential genes (Online GEne Essentiality (OGEE) database); knockdown of these genes causes lethal or infertile phenotypes in *Mus musculus* (methods). For all centrality metrics, gene centrality was proportional to biological essentiality (**Fig. 3B,C**), which supports the reliability of our networks and the validity of applying node centrality theories to single-cell data. *Pagerank* centrality was the best predictor of gene essentiality, and was the most consistent measure across organs (**Fig. 3D**). *Eigenvalue* was the worst predictor, its performance possibly depending on the network structure (**Fig. S5**). Next, to assess genes’ organ-specific centrality and how this relates to biological functions, we compared the centrality of genes across organs. In 11 regulatory networks we identified genes that were central for single or multiple mouse organs (**Fig. 3E, Fig. S2B**). Genes that were central in multiple organs appeared more essential than organ-specific genes (**Fig. 3G, Fig. S2C**). In line with this, shared central genes were associated with general housekeeping functions (e.g. gene expression or metabolic processes), whereas organ-specific central genes were associated with organ-specific processes (GO enrichment, **Fig. 3H**). Examples include *epidermis development* in the skin *(p<3.2×10^−5^), regulation of blood vessel diameter* in the heart (*p<4.1×10^−5^*) and *regulation of neuron apoptotic process* in the brain (*p<5.8×10^−4^).* Importantly, regulatory network analysis provided biologically relevant information not captured by gene expression levels alone. In fact, most organ-specific central genes are not up-regulated in their respective organs (**Fig. 3F**). This implies that gene expression levels are not an adequate measure of the importance of such genes for their underlying biological system. Overall, our framework for single-cell network analysis was capable of exposing functional regulatory structures and key genes that are undetectable by current computational strategies. We believe that this approach will be very valuable and broadly applicable for interpreting healthy and diseased complex biological systems. For the latter, regulatory networks will allow us to detect the molecular fingerprints of perturbations and to identify key driver genes for disease.

**Figure 3.**
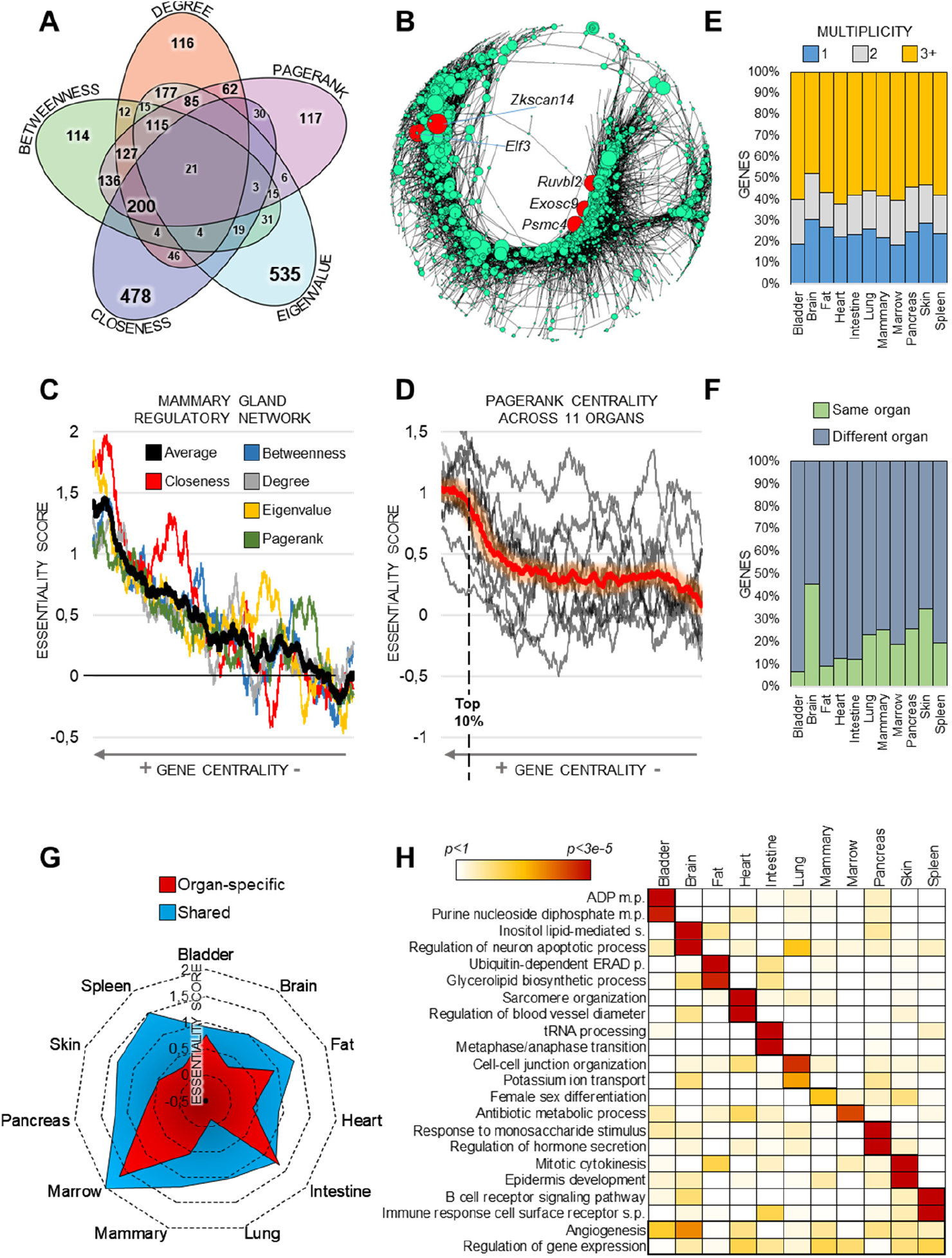
Regulatory networks inferred from 11 organs of the mouse body. (A) Marginal overlap of genes of different centralities in the brain network (top 20% genes). Additional organs in **Fig. S4**. (B) Visual representation the mammary gland network, in which node size is proportional to *pagerank* centrality. The top five central nodes (red color) are all genes classified by OGEE as biologically essential. (C) Genes (2403) of the mammary gland regulatory network sorted by centrality. Increasing centrality corresponds to higher biological essentiality for all centrality measures. (D) Genes sorted according to their *pagerank* centrality. High centrality (roughly top 10% of genes) corresponds to high biological essentiality. (E) The central genes (here, *pagerank*, top 20%) of each organ classified by their multiplicity. Multiplicity=1 means that they are central only in that organ, whereas multiplicity=2(3+) means that they are also central in additional organs (total of 2, or 3 or more, organs). Additional organs in **Fig. S2B**. (F) Analysis of genes which are i) central in at least one organ (*pagerank*) and ii) up-regulated in one organ compared to others. Intriguingly, most of the genes central in a given organ are expressed to a significantly higher extent (p<0.05) in a different organ. Additional organs in **Fig. S2D.** (G) Radar plot of the *pagerank* essentiality score for the central genes of each organ (top 20%), partitioned in either organ-specific or shared (multiplicity 3+). The latter have higher biological essentiality and reach a significant p-value for all organs (random permutations, p<0.005; **Fig. S2C**). (H) Heatmap of GO enrichments using *pagerank* centrality.

### Altered regulatory network architecture in pancreas from type 2 diabetes (T2D) patients

We considered that regulatory networks and gene centralities would be particularly informative about latent disease-related regulatory changes that are invisible to current analytical approaches. Thus, we generated healthy and T2D regulatory networks for 2,491 single-cell transcriptomes from diabetes patients and controls (Segerstolpe et al., 2016). First, we studied disease-related changes in *pagerank* centrality, a metric originally conceived to rank the popularity of websites. Nodes with high *pagerank* centrality indicate “popular” genes involved in multiple regulatory pathways. We hypothesized that genes with altered *pagerank* centrality would represent T2D regulatory changes with high functional impact on disease pathology. We found 162 genes with significantly increased *pagerank* centrality in T2D, despite showing equal expression levels (p>0.05) in T2D patients and healthy controls (**Fig. 4A,B**). In addition, we detected 10 genes, including *insulin* (*INS*), with increased *pagerank* that were significantly down-regulated in T2D (p<0.05). Consistent with known disease pathology, *insulin* was the most down-regulated gene (p<2.2×10-264), but had significantly higher pagerank centrality (from 0.3 to 0.9; Fig. 4A,B). This shows that insulin is a crucial limiting factor in the T2D network, and further emphasizes its pivotal role for the disease. Next, we used GO and GSEA to confirm the importance of the 172 genes with increased pagerank for pancreas function. Gene set enrichment supported their role in diabetes pathophysiology, as illustrated by the overrepresentation of terms such as “onset of diabetes in the young” signature (p<0.016; **Fig. 4C**). Genes showing changes in the remaining centralities (eigenvalues, closeness, betweenness, degree) were also enriched in diabetes-related functions, further highlighting the value of our method for interpreting scRNA-seq data (**Table S1**).

**Figure 4.**
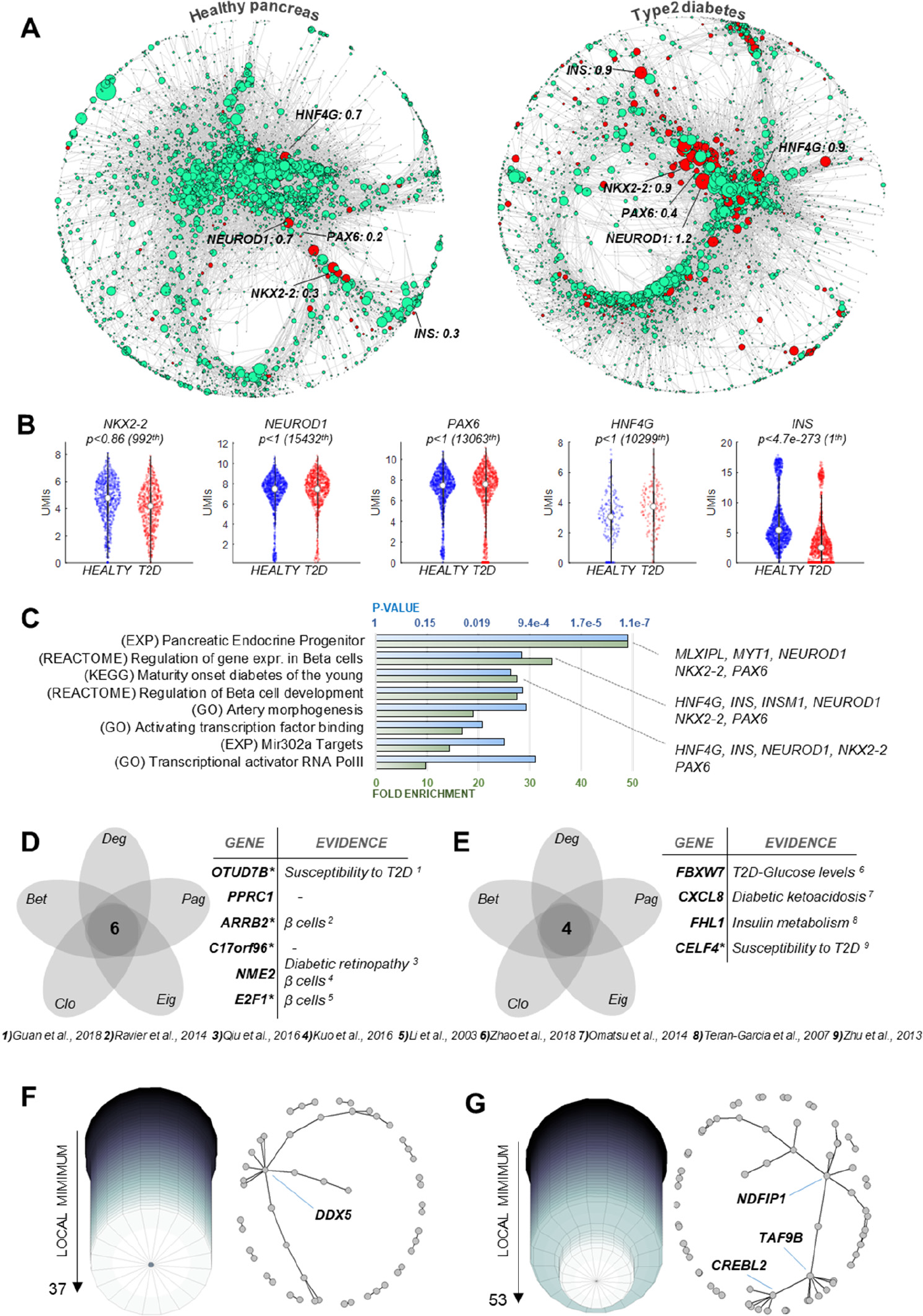
Changes in centralities and dynamical properties in the pancreas of type 2 diabetes (T2D). (A) Network from healthy and T2D subjects, in which node size is proportional to its *pagerank* centrality. In red, the nodes which are present in both networks and have higher *pagerank* in T2D (84 nodes). (B) Violin plots, p-values and ranking in differential expression in healthy vs. T2D tissue using all cells (1313 control cells vs 1178 T2D cells). (C) GO and GSEA enrichments showing overrepresentation of diabetes-related functions in nodes with increased *pagerank.* (D-E) Several of the genes showing simultaneous decrease (D) or increase (E) of the five centralities have been implicated in diabetes. (F-G) Degree of the monotonicity (residual negative edges) was 37 for control (F) and 53 for T2D (G). Nodes carrying residual negative edges are highlighted in the networks plots.

Finally, we identified genes with a simultaneous increase or decrease in the five centralities, which we expect to drive essential regulatory changes in T2D. We detected 4 (6) genes with repeated increased (decreased) centrality, most of which have previously been linked to diabetes pathology (**Fig. 4D,E**) (Guan et al., 2018; Kuo et al., 2016; Li et al., 2003; Omatsu et al., 2014; Qiu et al., 2016; Teran-Garcia et al., 2007; Zhao et al., 2018; Zhu et al., 2013). For example, *ARRB2*, a gene with a demonstrated role in *β* cell development (Ravier et al., 2014), showed no differential expression (p>0.05) but was simultaneously decreased in all six centrality measures. This is particularly remarkable because *β* cells were the most deregulated cell type in the original analysis (Segerstolpe et al., 2016), which, however did not detect the importance of *ARRB2* this context. This further supports the notion that generating global regulatory networks from single-cell data provides important insights into the pathological mechanisms of diseases.

### Monotone behavior of healthy and diseased pancreatic tissue

Dynamical behavior is perhaps the most important aspect of biological modeling, and describes how systems respond to input or perturbation. Biological regulatory networks have been suggested to display nearly monotone behaviors (Iacono and Altafini, 2010; Iacono et al., 2010), i.e. the prevalence of predictable, bounded trajectories over “chaotic”, oscillatory behaviors. Generally, abundant negative signs (negative correlations between genes) favor non-monotone behaviors. A social network is an intuitive parallel of this, in that unfriendly relationships (negative signs) increase social tension and decrease social monotonicity (Facchetti et al., 2011). We considered it interesting to assess whether our scRNA-seq-derived regulatory networks preserve the near-to-monotone behavior previously found in manually curated networks (Iacono and Altafini, 2010; Soranzo et al., 2012). The T2D regulatory network possessed more negative edges than the healthy counterpart, suggesting that diabetes causes an increase in chaotic signaling in the pancreas.

Computing the distance to monotonicity (how many negative edges must be removed to achieve monotonicity) of large networks is a complex, NP-hard (non-deterministic polynomial-time) problem, for which there are only approximate solutions. We used a greedy heuristic based on gauge transformations and previously shown to be the most accurate solution for large networks (Iacono et al., 2010). As expected from the number of negative edges, the T2D network was less monotone than the healthy network (53/52035=0.101% compared to 37/48549=0.076% residual negative edges, respectively) (**Fig. 4F,G**). The nodes carrying these residual negative signs have potentially chaotic effects in the dynamic behavior of the network. Herein, we identified these potentially chaotic transcription regulators in healthy (*DDX5*) and T2D (*NDFIP1, TAF9B* and *CREBL2*) networks (**Fig. 4F,G**). However, both networks were extremely close to monotone, in comparison to previously reported regulatory networks (Yeast 3.8%, E.coli 11.2% residual negative edges) (Iacono et al., 2010). Hence, the decrease in monotonicity in T2D was not sufficient to suggest that diabetes induces “chaotic” signaling in the pancreas. Nevertheless, this approach illustrates the potential for going beyond clustering-based phenotype analysis of single cells, allowing us to infer advanced system properties, such as dynamical behaviors of organs in health and disease. While we did not observe increased “chaotic” signaling in T2D, is it still intriguing to speculate that this is the case in other diseases.

### Network-driven interpretation of differentially expressed genes

Differential expression (DE) is the backbone of most analytical pipelines for RNA-seq. A typical challenge is to interpret differentially expressed genes and identify functionally important events. This is generally achieved by i) focusing on the genes with most significant p-value, ii) integrating external databases (GO or GSEA) to elucidate key genes and pathways, or iii) using personal knowledge to identify previously annotated genes. However, none of these approaches guarantees an unbiased classification of biological importance. In fact, in DE analysis p-values rank genes by technical reproducibility, not by biological importance, and both external databases and personal knowledge can be biased. Single-cell regulatory networks can be used to provide an unbiased, hypothesis-free classification of the biological importance of genes, allowing us to automatically identify pivotal deregulated genes, which greatly facilitates data interpretation. Comparing gene expression in *β* cells between healthy and T2D individuals, we detected 911 genes up-regulated in T2D *β* cells (p<0.05; **Fig. 5A**). Ranking these genes by centrality rather than p-values (i.e. Z-scores) provided quantitative sorting by biological importance, allowing us to immediately focus on the most relevant candidates. For example, *NEUROD1* and *RCAN1* showed the highest centrality of all deregulated genes according to multiple metrics (**Fig. 5A,B**), suggesting that they are the most informative and biologically relevant. Interestingly, mutations in *NEUROD1* were associated with T2D (Malecki et al., 1999), whereas up-regulation of *RCAN1* was shown to cause hyperinsulinemia, *β* cell dysfunction and diabetes (Peiris et al., 2012). Notably, neither of these genes was highlighted with DE p-values (*NEUROD1* 2829th, p<2×10^−4^; *RCAN1* 4331th, p<0.05). This example highlights the high additive value of using single-cell regulatory networks and related node centralities to aid interpretation of DE results.

**Figure 5.**
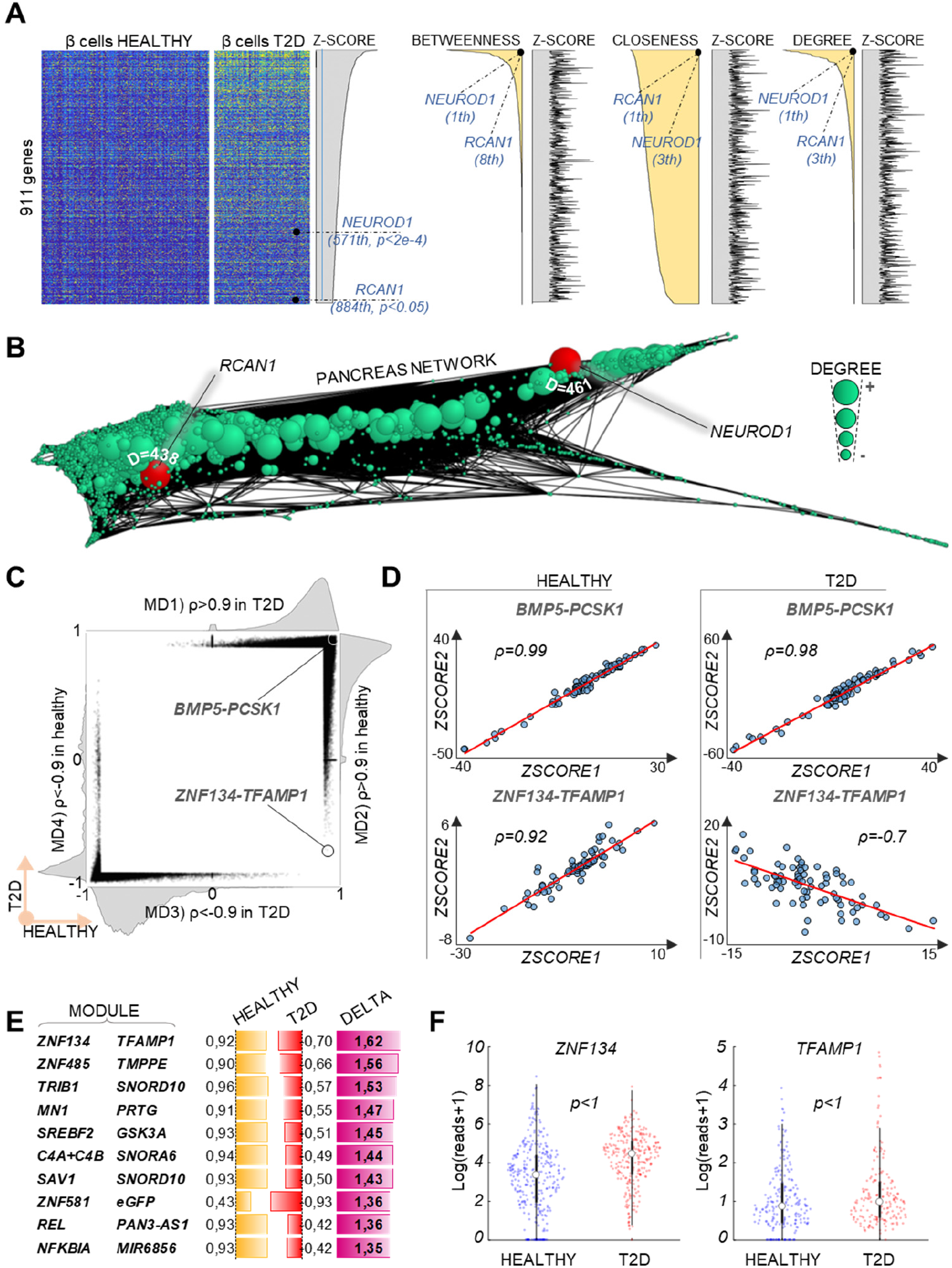
Prediction of gene importance in DE data and directional changes in correlations. (A) Heatmap of normalized expression values of 911 genes found significantly up-regulated in T2D β cells compared to healthy β cells (p<0.05, *bigSCale*) sorted by decreasing Z-score (i.e. Increasing p-values) or decreasing centralities (*betweenness, closeness* and *degree*). Biological importance of *NEUROD1* and *RCAN1* is highlighted by their high centrality, but not by their DE Z-scores. More in general, correlation between DE Z-scores and centrality is marginal, as shown by the erratic area plots of Z-scores sorted by centrality. (B) Pancreas regulatory network generated using all cells (both healthy and T2D), node size proportional to its degree. *NEUROD1* and *RCAN1* have high degree centrality (461 and 438, respectively). (C) Scatter plot and marginal distributions (MD1-4) of all pairwise gene correlations with |ρ|>0.9 (329’046 couples). The strong bias in the marginal distributions (especially MD1, 2, 4) indicates an overall similarity of correlations between healthy and T2D pancreas. (D) An example of a highly conserved correlation (*BMP5-PCSK1*) against the strongest inversion (*ZNF134-TFAMP1*). (E) Overview of top 10 inversions of correlations. F) Neither ZNF134 nor TFAMP1 display a significant change of expression (*bigSCale* DE).

### Inversions of gene correlations in T2D

Regulatory networks can be further interrogated to detect changes in local interactions, namely pairwise correlations between genes. With the rationale that gene pairs with directional changes in correlation represent rewired functional modules with pathological implications, we performed comparative analysis of the healthy and T2D networks. Overall, all pairwise correlations were highly similar (example of *BMP5* and *PCSK1*, **Fig. 5C,D**) under healthy and T2D conditions, which is remarkable considering that the data come from different donors and are subject to inter-individual variability as well as several confounding factors (e.g. age and weight). This indicates that our approach works transversely to confounding variables, ultimately exposing the true functional correlations between genes. Closer inspection revealed a number of modules (14) with strongly (ρ>1) inverted correlations, the most striking example being *ZNF134* and *TFAMP1*, which switch from a strong positive correlation in the healthy pancreas (ρ=0.92) to a negative correlation in T2D (ρ=–0.7). Neither of these genes showed a change in expression between conditions (healthy/T2D), which renders their altered functionality invisible to standard methods (**Fig. 5E,F**).

Several other genes displaying inverted correlations have previously been linked to diabetes, either by functional studies (*TRIB1*, glucose metabolism; *NFKBIA*, insulin resistance pathway) or as candidate disease genes in GWAS or gene expression studies (*TMPPE, PRTG* and *ZNF319*) (Ishizuka et al., 2014; Itani et al., 2002; Miller et al., 2010; Wanic et al., 2013; Yang et al., 2016; Zhang et al., 2018). Functionally, the most interesting are *SREBP2* and *GSK3A*, which have a direct mechanistic relationship and are both implicated in T2D, and which also switched from a positive to a negative correlation. SREBP transcription factors are major players in lipid metabolism and possibly insulin resistance, whereas GSK3 phosphorylates SREBP in the absence of insulin and AKT signaling, leading to its degradation (Henriksen and Dokken, 2006; Musso et al., 2013; Shao and Espenshade, 2012). Consequently, we can speculate that the reversal in correlations inferred from single-cell data is directly related to a change in insulin signaling and the degradation of SREBP2 through GSK3A.

In summary, the comparative analysis of single-cell-driven correlations is a suitable novel approach for disentangling the molecular mechanisms of diabetes, and further enlarges the repertoire of single-cell data analysis strategies available for meaningful data interpretation.

### Rewiring of microglia gene regulation in Alzheimer’s disease (AD)

To further evaluate the applicability of our network-based approach in a different disease context, we analyzed scRNA-seq data from immune cells (CD45+) derived from 5XFAD transgenic mice, a commonly used model for AD (Keren-Shaul et al., 2017). The dataset contains transcriptomes from 22,951 single cells from different disease stages (1-8 months, control and 5XFAD) as well as *Trem2^+/+^* and *Trem2^−/−^* AD and control mice (*Trem2* is a key receptor that modulates immune response). The variety of conditions makes the dataset particularly suitable for confirming the benefits of our unifying approach. In fact, instead of progressively fragmenting cells and conditions into stratified groups and clusters, we use all of the data as an input to generate regulatory networks for controls and AD. Overall, while the two networks were of comparable size (**Table 1**), we observed a general loss of connectivity in AD, which increased network sparseness and signal travelling time (shortest paths, **Table 1**). Consequently, centralities of several genes were different in the AD network compared to the control (**Fig. 6A**). Altered centralities were associated with different GSEA enrichments, further reflecting the fact that each centrality highlights different functional aspects. For example, genes that lost influence (*eigenvalue*) significantly overlapped with genes that were dependent on *Trem1* in monocytes (p<6.5×10^−6^). This observation is intriguing given the relevance of *Trem2* in the mediation of immune response in AD brains.

**Figure 6.**
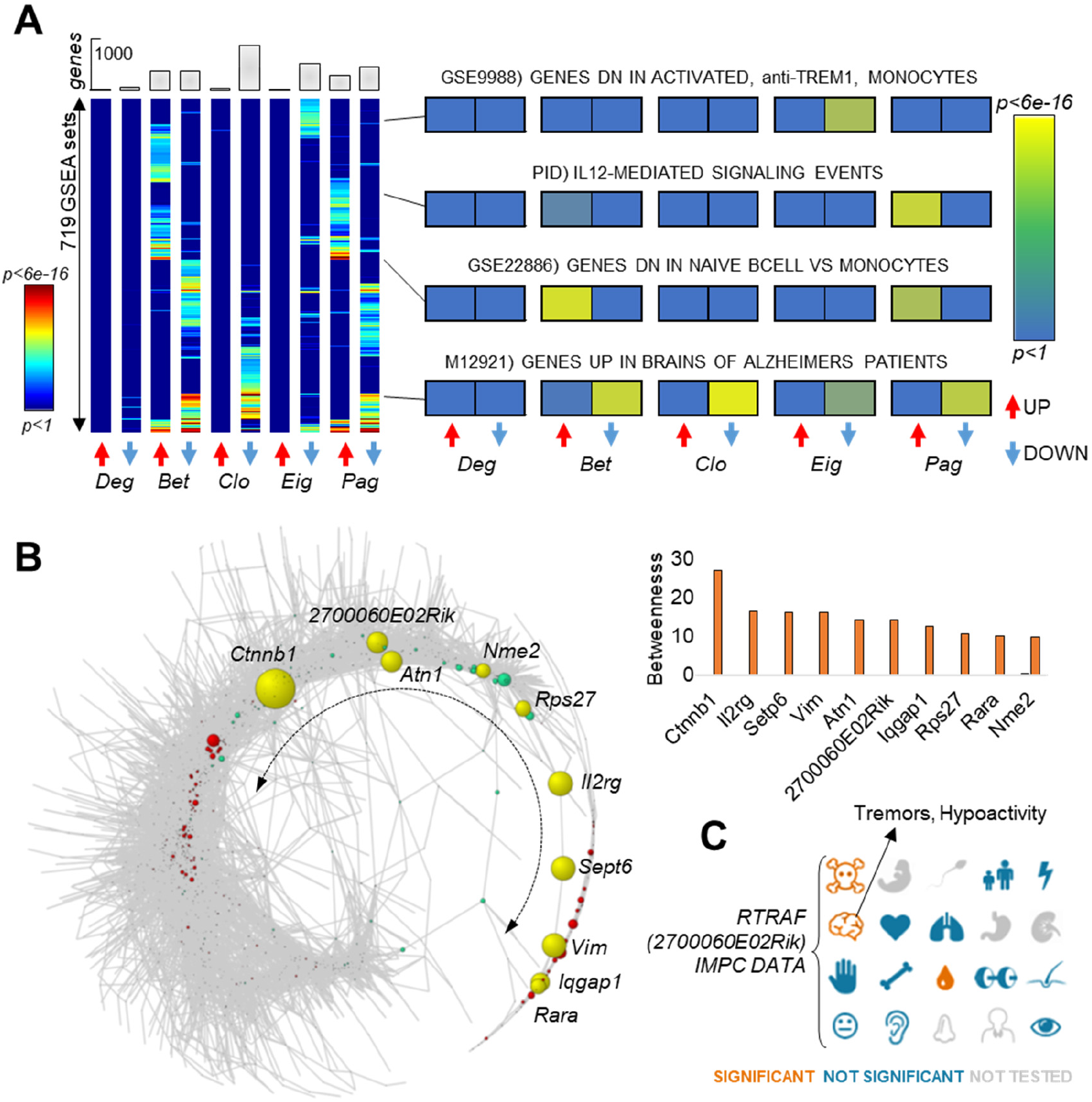
Rewiring of regulatory network in Alzheimer’s disease microglia introduces information bottlenecks. (A) GSEA enrichments for 10 lists of genes with altered centrality in AD compared to control network. For 5 centralities (up or down) 719 GSEA terms were found to be enriched with p<0.05 in at least one entry). The results show that the tested centralities provide insights into different functional pathways. (B) The AD regulatory network follows a circular shape in which the nodes along the terminal tail (yellow genes) become bottlenecks, i.e. acquire increased *betweenness.* The exact increase in *betweenness* is represented in the histogram. (C) International Mouse Phenotyping Consortium data for the transcript *2700060E02Rik*, alias *Rtraf.* Homozygous *2700060E02Rik* knockout is associated with embryonic lethality prior to tooth bud stage and heterozygous 2700060E02Rik knockout is associated with tremors, hypoactivity and increased eosinophil cell number.

Other function-centrality associations include genes that are up-regulated in AD patients (p<5.0×10^−12^), the Interleukin 12 signaling cascade (p<1.1×10^−8^) and genes that are down-regulated in naive B-cells compared to monocytes (p<3.5×10^−10^; **Fig. 6A**). Overall, *betweenness* showed the most dramatic changes of all centralities. In fact, the AD network was rewired into a circular shape, which in turn causes a number of genes to become information bottlenecks (**Fig. 6B**). Interestingly, *beta catenin 1* (*Ctnnb1*), part of the main pathway that regulates the onset and progression of AD (Ghanevati and Miller, 2005), showed the largest increase in *betweenness* (from 0.0% to 27.3%, **Fig. 6B**) and became the main bottleneck in the AD network. Among the top-10 genes with increased *betweenness*, we also found a poorly annotated transcript *2700060E02Rik* (Fig. 6C), whose heterozygous deletion was previously associated with tremors and hypoactivity, a common symptom of AD (Mouse Phenotyping Consortium, hwww.mousephenotype.org).

## DISCUSSION

During the last decade, single-cell transcriptomics has becoming increasingly important for deconvoluting the cellular architecture of complex tissues, and for classifying cells with categorizing principles. A holistic scenario, where single cells are combined to infer global regulatory networks, has not yet been comprehensively explored. There have been isolated studies using small-scale single-cell data to derive partial regulatory networks, although their reliability has been questioned (Chen and Mar, 2018). Hence, it remained unclear whether single-cell datasets can be analyzed using strategies other than clustering-based phenotyping.

The main obstacles that impede network analysis of single-cell data are the technical limitations inherent to the technology and the very large data volumes. Guo and coworkers used least square fitting on expression data from 28 epithelial cells and inferred a partial regulatory network of few hundred nodes and edges (Guo et al., 2015), however, without validating it. Further, the use of least square fitting is known to perform poorly with the sparse and low complexity of single-cell data (Fiers et al., 2018). In another work (Lim et al., 2016), 92 cells were analyzed using an asynchronous Boolean approach to refine literature curated models of hematopoiesis. Boolean approaches are not easily scalable (Fiers et al., 2018), and can therefore only be used to inspect reduced, specific sub-networks. Other studies applied graphical approaches not scalable to large-scale sequencing data (Chan et al., 2017; Hamey et al., 2017; Matsumoto et al., 2017; Moignard et al., 2013; Pina et al., 2015; Sanchez-Castillo et al., 2018; Wei et al., 2017), metrics not tailored to scRNA-seq specific features (Chan et al., 2017; Hamey et al., 2017; Moignard et al., 2013; Papili Gao et al., 2017; Pina et al., 2015; Wei et al., 2017), or dynamic process-specific approaches (Matsumoto et al., 2017; Moignard et al., 2013; Papili Gao et al., 2017). More recently, Aibar and colleagues developed the first tool designed to infer transcription factors and their target genes from single-cell data (Aibar et al., 2017). However, this tool ultimately applies clustering and phenotyping, without generating global regulatory networks.

In this work, we conceived an analytical framework for inferring large-scale regulatory networks from single-cell data. To confirm the viability of this approach, we generated a large and diverse repertoire of regulatory networks in healthy and diseased contexts. To support network interpretation, we applied advanced tools from graph theory, and validated this strategy thoroughly at multiple levels. Importantly, we showed that regulatory networks derived from single-cell data can be used to obtain novel and biologically relevant insights into the molecular architecture of complex systems and the pathophysiology of diseases. This work represents an important leap forward in the field of single-cell analysis for the reasons described below.

*First*, we conducted the first large-scale analysis of global regulatory networks using single cells. We processed datasets from up to 8,000 single cells into networks with up to 60,000 edges and 7,000 nodes, going far beyond previous studies (Guo et al., 2015; Lim et al., 2016).

*Second*, we conceived a metric which consistently identified hidden correlations within the single-cell dataset. The metric was developed within the framework of *bigSCale* (Iacono et al., 2018), and was specifically tailored to single-cell data, diminishing the effect of data sparsity, confounding factors, and other technical artifacts. Thereby, it removes main obstacle to processing scRNA-seq data into regulatory networks.

*Third*, we studied the global and local properties of networks using advanced tools from graph theory, enabling a comprehensive characterization. Some of the concepts, such as monotonicity, have not yet been applied to data-driven regulatory networks, but have proven to be extremely powerful for understanding the biological systems analyzed here.

*Fourth*, we validated our results at multiple levels. Specifically, we validated inferred correlations between transcription regulators and target genes via experimental signatures of perturbed biological systems. In line with previous evidence, the inferred networks were scale-free (Albert, 2005; Balaji et al., 2006; Noort et al., 2004). The centrality of genes was validated using external experimental datasets of essential genes (OGEE database), supporting their biological relevance. Further, we validated the functionality of organ-specific central genes in their respective tissue contexts (GO enrichment). Lastly, we found that genes with altered centrality in T2D and AD strongly overlap with previous known disease mechanisms.

*Fifth*, we compared the results from the regulatory network approach with those from conventional DE analysis. Notably, we found that networks repeatedly disclosed latent variation and features that were invisible to standard analysis. Moreover, we showed that gene centrality analysis was able to work in *synergy* with differential expression analysis to provide an unbiased, quantitative ranking of biological importance from dysregulated genes. To our knowledge, this is a unique strategy for deducing a data-driven biological ranking without the need to incorporate external information (e.g. GO or GSEA) or personal knowledge.

*Sixth*, we have completed the first single-cell, network-driven analysis of diseased samples. Here, graph-based tools allowed us to enhance our understanding of their molecular pathology. In general, given its holistic rather than classifying use of single cells, we propose that the network approach is particularly well-suited for complex experimental designs with multiple confounding factors, such as clinical samples displaying sex-, age- and weight-related differences.

In summary, we have shown that regulatory network approaches can be applied to large-scale single-cell datasets, and can be used to maximize the biologically relevant information obtained. Testing consistency across multiple networks will allow us to determine the completeness of the captured regulatory interactions, and this should be the primary future task.

## METHODS

### Inferring gene expression correlations and regulatory networks from scRNA-seq data

Single-cell sequencing is characterized by a series of technical limitations that generate artifacts, such as drop-out events, irregular sequencing depth and low library complexity. First, Drop-out events represent expressed genes that are undetected by scRNA-seq for technical reasons, resulting in zero values in the expression count matrix. These events make single-cell datasets considerably sparser than bulk RNA-seq datasets. Drop-out events are perhaps the most important factor affecting the performance of correlation methods, such as *Spearman* or *Pearson*, applied directly to expression count data. Second, irregular sequencing depth is caused by the uneven (non-normalized) loading of single-cell libraries into the sequencing reaction. Consequently, we observe large fluctuations in sequencing depth between cells, which can only partially be addressed by data normalization. Third, single-cell data present a reduced dynamic range of expression values, which is a further challenge for the performance of correlation methods. In fact, as it is not possible to entirely remove the effects of read distribution biases, traditional correlation metrics have suboptimal performance with this data type. Since these technical artifacts concur to mask correlations when using expression counts, we envisaged that a change of variable would greatly improve the performance of the correlation methods, thereby allowing us to infer the regulatory networks. To this end, we devised the following steps:

#### 1) Data pre-processing

Datasets were analyzed using the *bigSCale* framework, which handles the noise and sparsity of scRNA-seq data using an accurate numerical model of noise. The framework includes modules for differential expression analysis and unsupervised cell clustering. All datasets were processed using *bigSCale* under default parameters, with the exception of parameters regulating the granularity of clusters. *BigSCale* was set to the highest granularity in order to produce the highest number of clusters, the rationale being to segregate cell subtypes and subtle cell states, so as to improve the resolution and quality of inferred correlations.

#### 2) Measuring correlations in the Z-score space

After clustering the cells to highest feasible granularity, we used *bigSCale* to run an iterative differential expression (DE) analysis between all pairs of clusters. For *x* clusters, this results in a total of *x*(x-1)/2* unique comparisons, each yielding a Z-score for each gene that indicates the likelihood of an expression change between two clusters. This allows us to compute correlations between genes using Z-scores instead of expression values. For correlation analysis, we used *Pearson, Spearman* and *Cosine* metrics. We also tested the mutual information to detect non-linear correlations. However, in the Z-score space this resulted in an excessive number of false positives. Specifically, mutual information repeatedly identified significant dependencies for which one of the two variables was linearly independent of the other (slope=0). Nevertheless, linear correlations in the Z-score space can also reflect non-linear correlations in the original expression space. Hence, we chose to rely exclusively on a solid measure of linear correlation in the Z-score space via *Pearson, Spearman* and *Cosine* metrics. The final correlation for each pair of genes was computed as the lowest (worst) between *Pearson* and *Cosine* (*Spearman* is used in a later stage as a further control).

#### 3) Building a regulatory network

In the next step, we retained significant correlations to define the edges of the regulatory network. Notably, the distribution of correlations is influenced by biological and technical factors. For example, increased cell numbers or sequencing depth results in a higher number of significant correlations. Consequently, to compare regulatory networks inferred from different datasets, we must first adjust for technical factors, for which we used an adaptive rather than fixed correlation threshold. Specifically, the inferred networks were built by retaining the top 0.1% correlations. Using this relative correlation threshold prevents technical factors from producing artificial differences when comparing different networks. Although relative thresholds could result in the inclusion of non-significant correlations (e.g. ρ=0.4), we did not observe such events in any of the inferred networks, with most adaptive thresholds set between ρ_thresh_=0.9 and ρ_thresh_=0.99. The lowest (worst) adaptive threshold was ρ_thresh_=0.84 for the AD network, which is still significant. *Spearman* correlation is used as a further control to discard weak correlations. Specifically, final correlations for which |ρ_spearman_|<|ρ_thresh_-0.15| were considered null.

In a final step, the undirected network is polished to retain only the edges that represent actual regulatory links. To this end, we utilized GO annotations (version 24/03/2017) to extract putative regulators of transcription (GO:0010468 “regulation of gene expression”). We discarded from the network edges representing pairs of genes of which neither was annotated as “regulator of gene expression”, as we considered these to be spurious co-expression links. Alternatively, more specific GO terms could be used for network polishing (e.g. GO:0006355 “regulation of transcription, DNA-templated” or GO:000370 “DNA-binding transcription factor activity”). However, we opted for a broader term so as to include in our networks all possible regulatory layers, including indirect signaling events.

### Validation of network edges with external datasets

Our inferred regulatory links represent putative events of transcriptional regulation on target gene(s). We chose to also include indirect regulation events that do not imply the direct binding of a transcription factor to the promoter of the target gene(s). By filtering the edges using the broad GO term “regulators of transcription”, we included all possible regulatory layers, including transcription co-factors, epigenetic mechanisms, regulation of RNA stability/degradation, and signaling cascades. Consequently, neighboring genes (genes connected by an edge) are likely to belong to a common pathway and should be similarly affected when the system is perturbed. GSEA contains an extensive collection of experimental signatures associated with perturbation of biological systems, which we used to independently validate each edge in our networks. To detect significant enrichment of co-occurrences, we applied the Fisher’s exact test. Edges with significant p-values imply that the related genes are activated/deactivated together in experimentally perturbed systems significantly more often than expected by chance. The distribution of edge-wise fold enrichment (i.e. how often the edges translate into cooccurrences in GSEA signatures) was biased towards positive values for all mouse organs tested, indicating an overall simultaneous modulation of neighboring genes (**Fig. 2E,F**). Co-regulation was further supported by significant p-values (e.g. **Fig. 2G, Fig. S2A**), especially when considering edges with higher numbers of associated GSEA signatures (small gene sets are less likely to yield significant p-values in the Fisher exact test). Notably, we inferred organ-specific regulations, whereas the GSEA signatures are collected from a highly heterogeneous set of biological sources. Inevitably, some of our organ-specific regulatory links will be not backed-up by GSEA signatures, which explains why we could not validate not all individual edges in our networks.

For all the GO (version 24/03/2017) and GSEA (version v6.0) enrichment analyses we used hypergeometric distribution with *Bonferroni* correction.

### Validation of network hubs with gene essentiality

To elucidate whether the hubs in our networks represent essential regulatory factors, we took advantage of the Online GEne Essentiality (OGEE) database. This database provides an unbiased, comprehensive catalogue of the essentiality of experimentally tested genes across species. In this setting, we used the *Mus musculus* dataset (available at http://ogee.medgenius.info/browse/Mus%20musculus), which lists the essentiality status for 9,402 mouse genes. To quantify the essentiality of each set of hubs, we computed an essentiality score (ES), as:

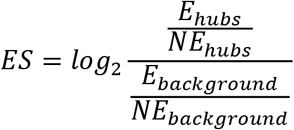

where *E_hubs_* and *NE_hubs_* are the number of essential and non-essential hubs, and *E_background_* and *NE_background_* are the number of essential and non-essential genes in the OGEE dataset, respectively.

To assess the significance of each ES, we computed the empirical probability of finding a score of the same magnitude by chance. Specifically, given a set with N hubs, we sampled N random genes from the OGEE dataset and calculated the ES. We repeated this process 10,000 times, and from the resulting distribution, we used the one-tailed p-value as the proportion of random ES that are equal to or greater than the observed ES. After calculating one p-value for each ES, we corrected for multiple testing by applying a Benjamini-Hochberg correction to the vector of p-values.

### Detection of changes in centralities

We evaluated two different approaches for ranking nodes according to their change in centrality. The first approach identifies the highest absolute change in centrality: Δ*c* = *c_a_ – c_b_*, where Δ*c* for each node is defined as the difference in its centrality between networks A (*c_a_*) and B (*c_b_*). Next, we selected the 1000 nodes with the greatest change in centrality (either positive or negative). The change in centrality was then integrated with the p-values of the DE analysis (*bigSCale*, standard parameters) to identify genes undetected by DE (**Figs. S6A**). In an alternative approach to identify relative changes in centrality, we searched for dispersed nodes lying outside the proportional relationship between *c_a_* and *c_b_*. We performed non-linear fitting (smoothing spline) to derive a confidence interval of the dispersion. Nodes that showed overdispersion at p<0.05 were defined as having altered centrality (**Fig. S6B**). Ultimately, we did not use this analysis in the manuscript, opting for the absolute change only (first approach). This is because relative changes in centrality, as measured by overdispersion, were biased towards small changes in centrality, which were important at a relative level, but irrelevant at the absolute level.

## ACKNOWLEDGMENT

HH is a Miguel Servet (CP14/00229) researcher funded by the Spanish Institute of Health Carlos III (ISCIII). This work has received funding from the Ministerio de Ciencia, InnovaciÓn y Universidades (SAF2017-89109-P; AEI/FEDER, UE). Core funding is from the ISCIII and the Generalitat de Catalunya.

## AUTHOR CONTRIBUTIONS

HH and GI conceived the study. GI developed *bigSCale* and performed the network and statistical analysis. RM conducted the network analysis of mouse organs. HH and GI wrote the manuscript. All authors read and approved the final manuscript.

---

**Fig. S1.**
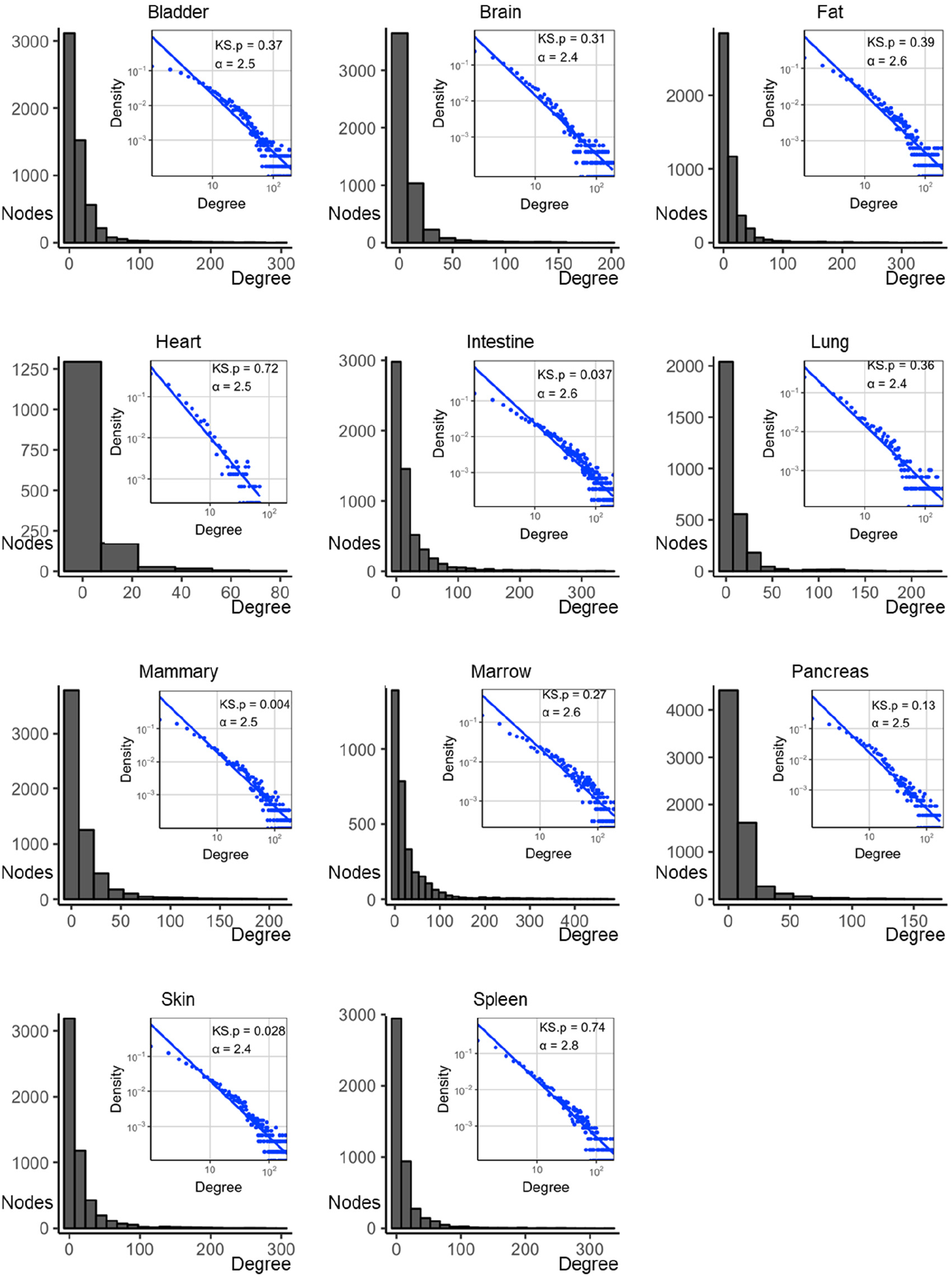
Single-cell gene regulatory networks are scale-free. The degree distribution of the single-cell gene regulatory networks derived for 11 organs shown in linear (histogram) and logarithmic scale (scatter plot). Each distribution was fitted to a power-law distribution, and the p-value of the Kolmogorov-Smirnov test (KS.p) and the degree exponent of the power-law (alpha) are shown for each network. All networks are scale-free (p>0.01) apart from the mammary gland which slightly deviates from the exact scale-free distribution (p<0.004).

**Fig. S2.**
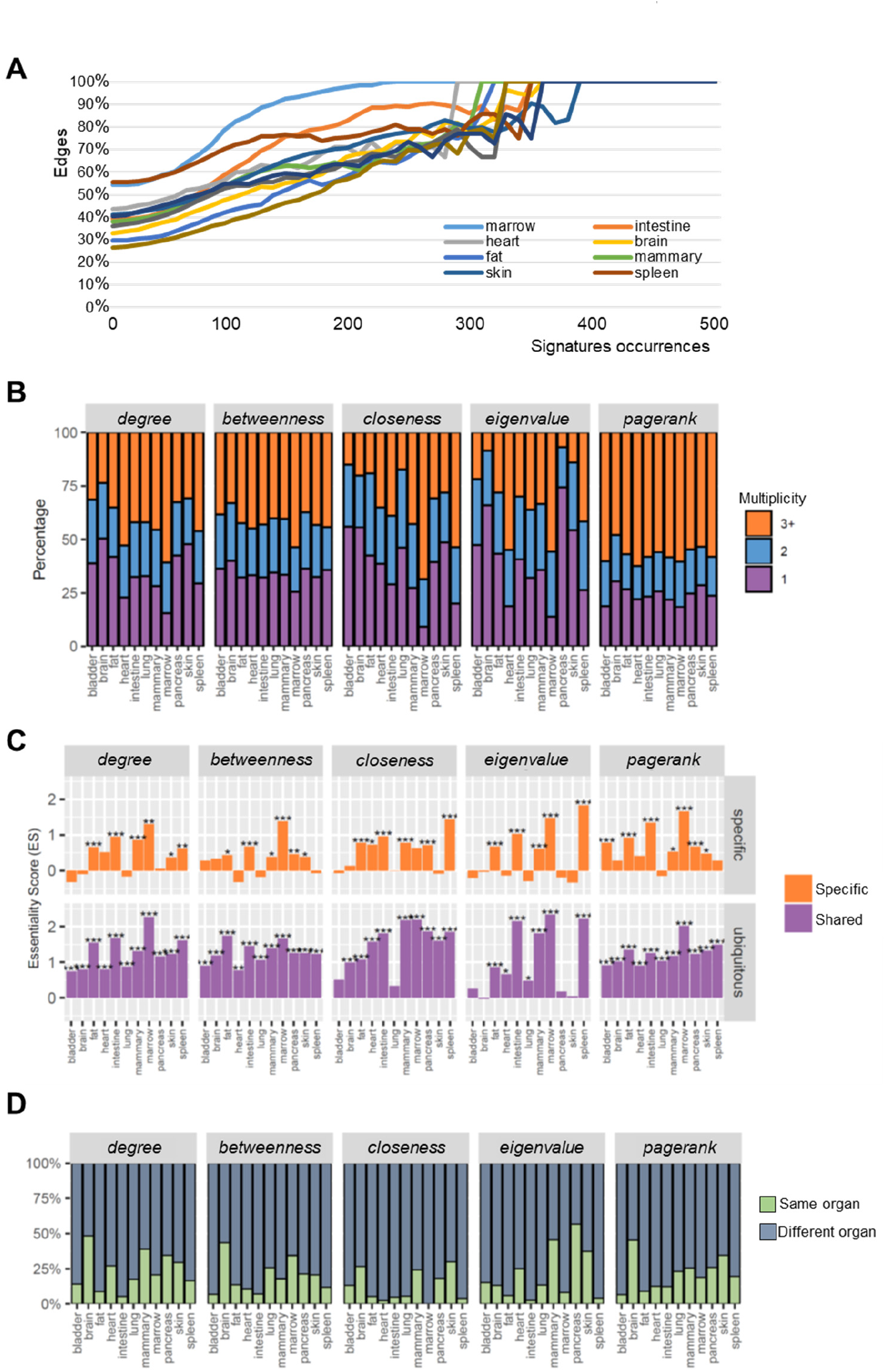
Validation of inferred networks and analysis of multiplicity. (A) Percentage of validated edges validated with *p<0.05* (y-axis) filtered by the number of GSEA occurrences (x-axis) for each organ. Increasing occurrences correspond to higher percentages of edges validated by significant p-values (p<0.05). (B) The central genes (top 20%) of each organ classified by their multiplicity for all tested centrality measures. Multiplicity=1 means that they are central only in that organ, whereas multiplicity=2(3+) means that they are central also in additional organs (total of 2, or 3 or more, organs). (C) Comparison of ES score of organ-specific central genes (multiplicity=1) against shared central genes (multiplicity 3+). The latter have higher biological essentiality (* p<0.05, ** p< 0.01, *** p<0.005, random permutations, see methods). (D) Analysis of genes which are i) central in at least one organ and ii) up-regulated in one organ compared to others. Intriguingly, most of the genes central in a given organ are actually expressed to a significantly higher extent (p<0.05) in a different organ. The brain has the highest amount of genes which are central and more expressed (compare to other organs) at the same time.

**Fig S3.**
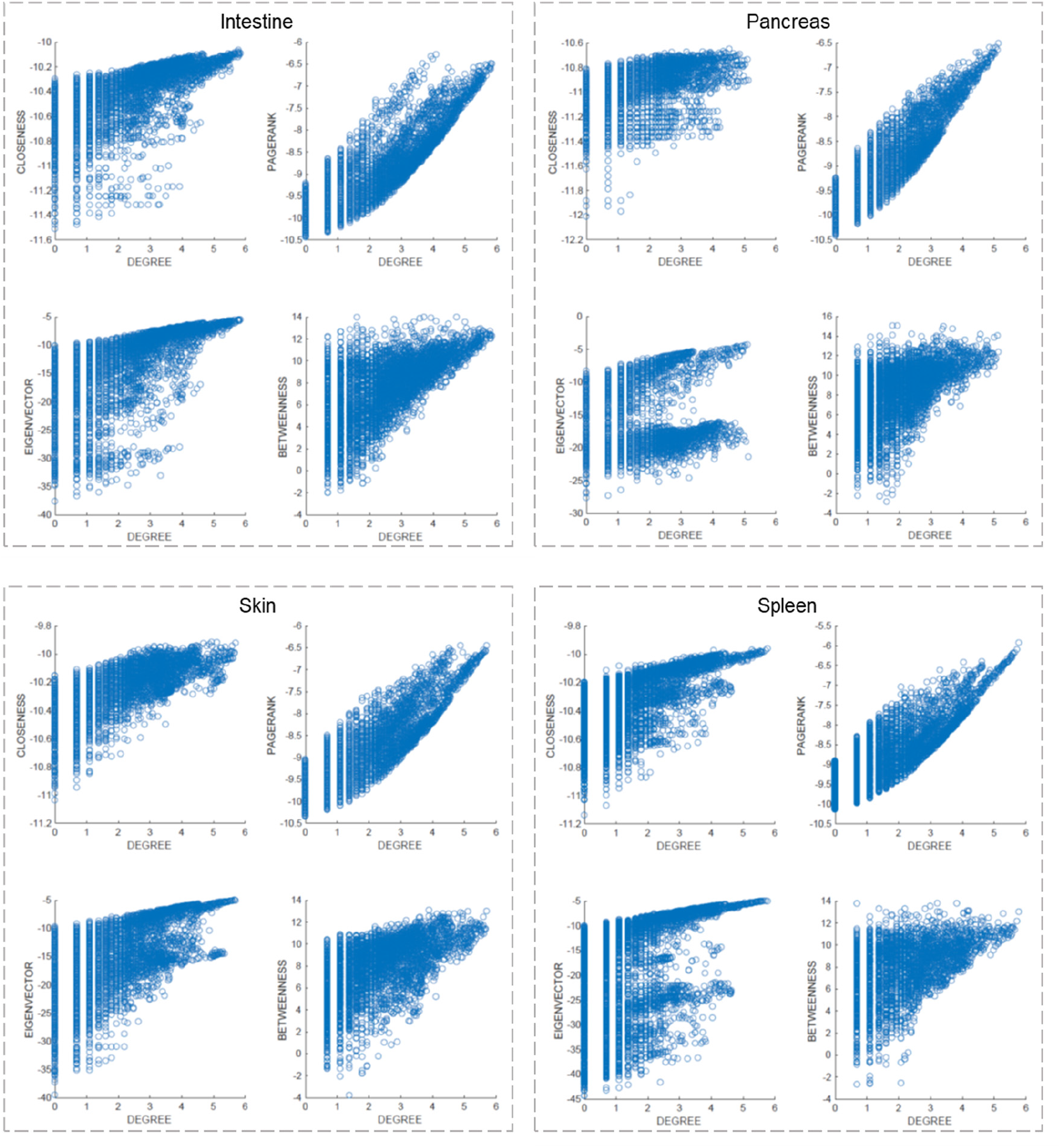
Relationship between degree and other centralities. Scatter plots between *degree* (log-scale) and the other centralities. Degree is perhaps the most simple and direct measure of centrality. Nodes with high degree have many connections and are therefore more likely to be central also in the other metrics. In line, degree and the other metrics show a general positive correlation, as shown in the examples of the intestine, the pancreas, the skin and the spleen. However, the other metrics are able to capture types of node importance (i.e. centrality) which the degree cannot. This is shown by the sparseness and/or multimodality of several distributions, such as for example *degree* and *pagerank* in the intestine, where nodes with the same degree can present widely different *pagerank* centrality.

**Fig S4.**
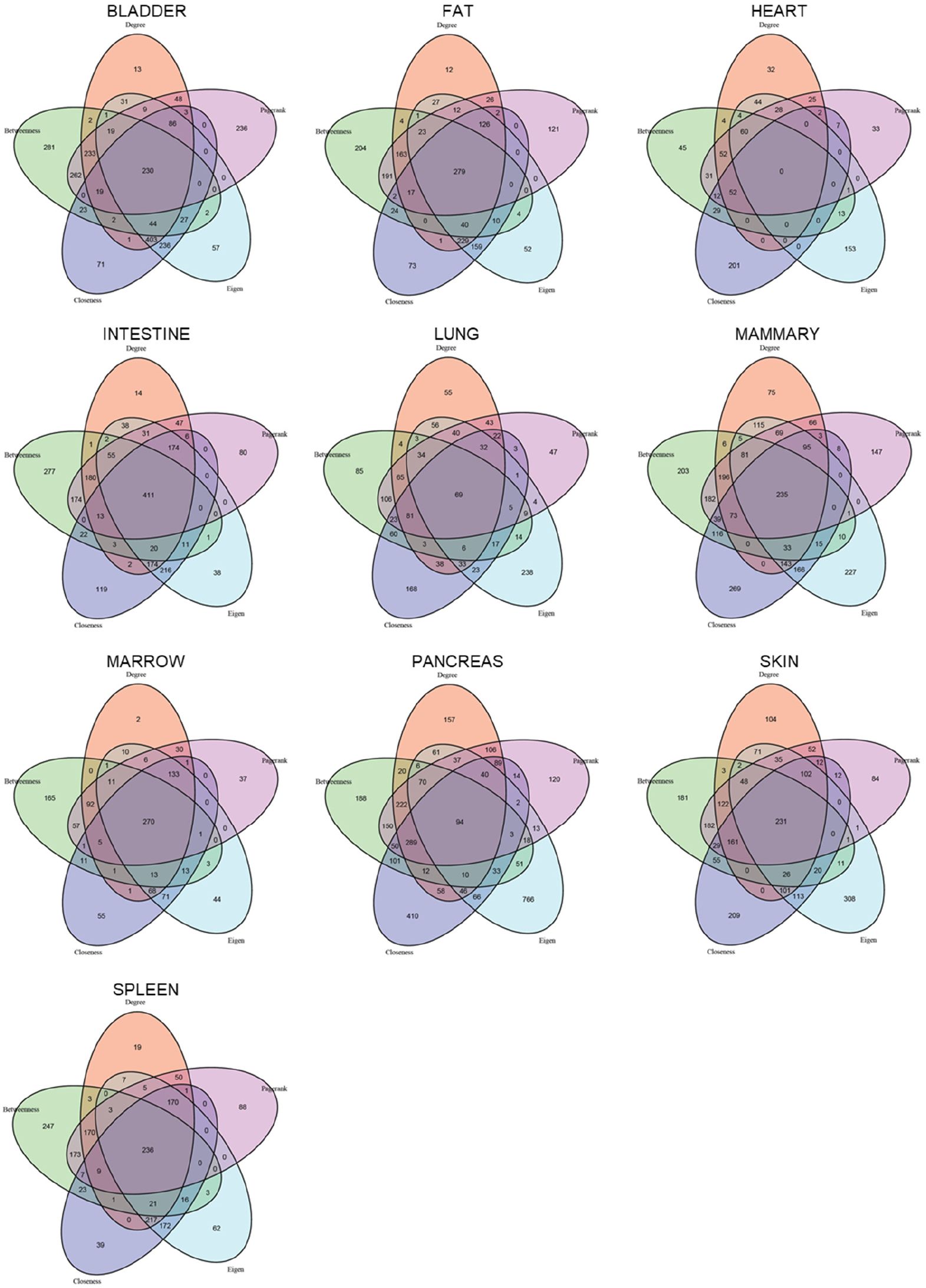
The central genes of different metrics show marginal overlap. Venn diagrams intersecting the genes central (top 20%) in different measures.

**Fig S5.**
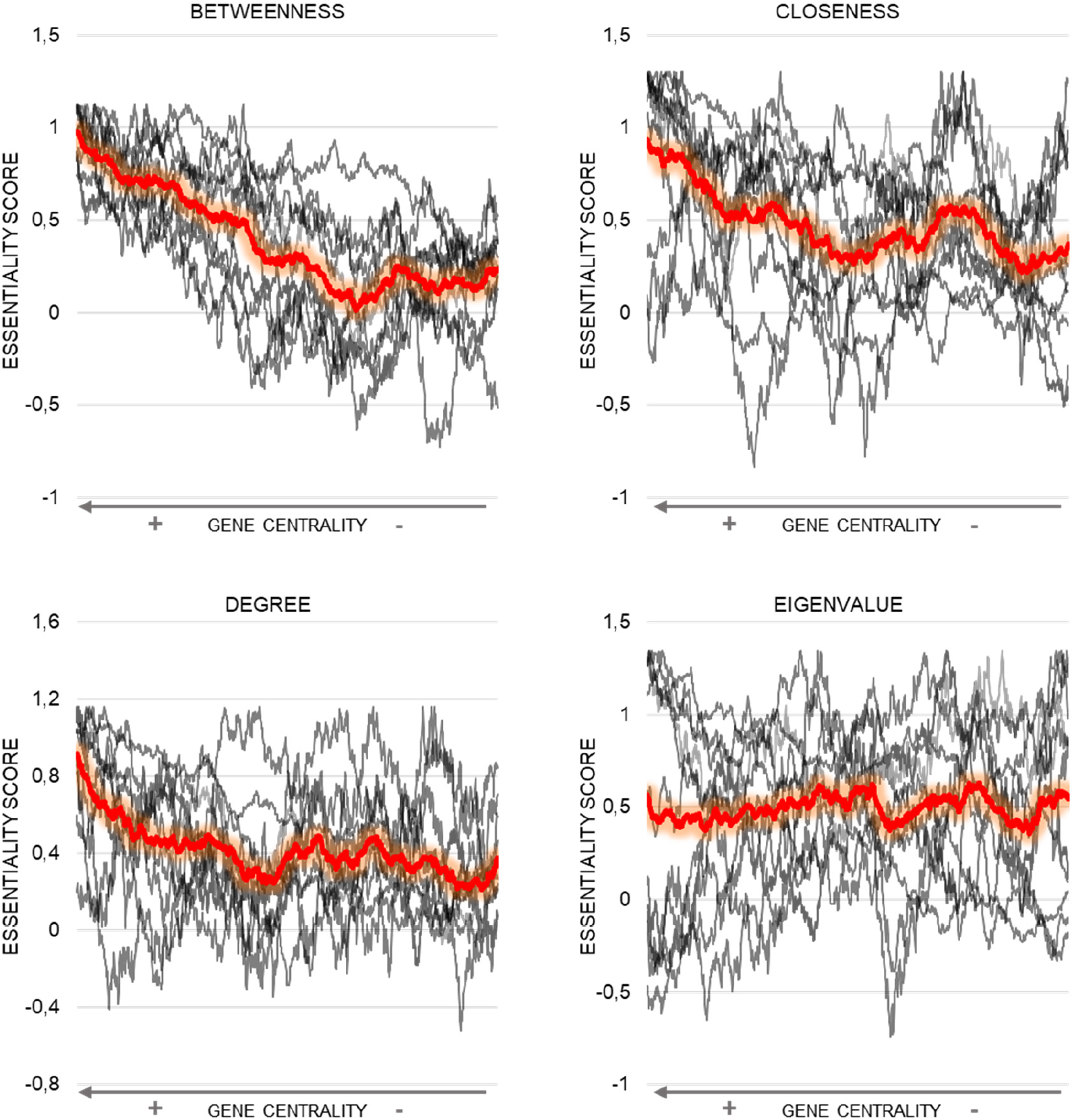
Relationship between gene centrality and biological essentiality. Genes sorted according to their centrality. The top central genes (left side of the x-axis) in *betweenness, closeness* and *degree* show the highest biological essentiality (ES score, OGEE database, see methods). *Eigenvalue* has an unstable performance (working for some organs and not for others), possibly depending on the structure of the network.

**Fig S6.**
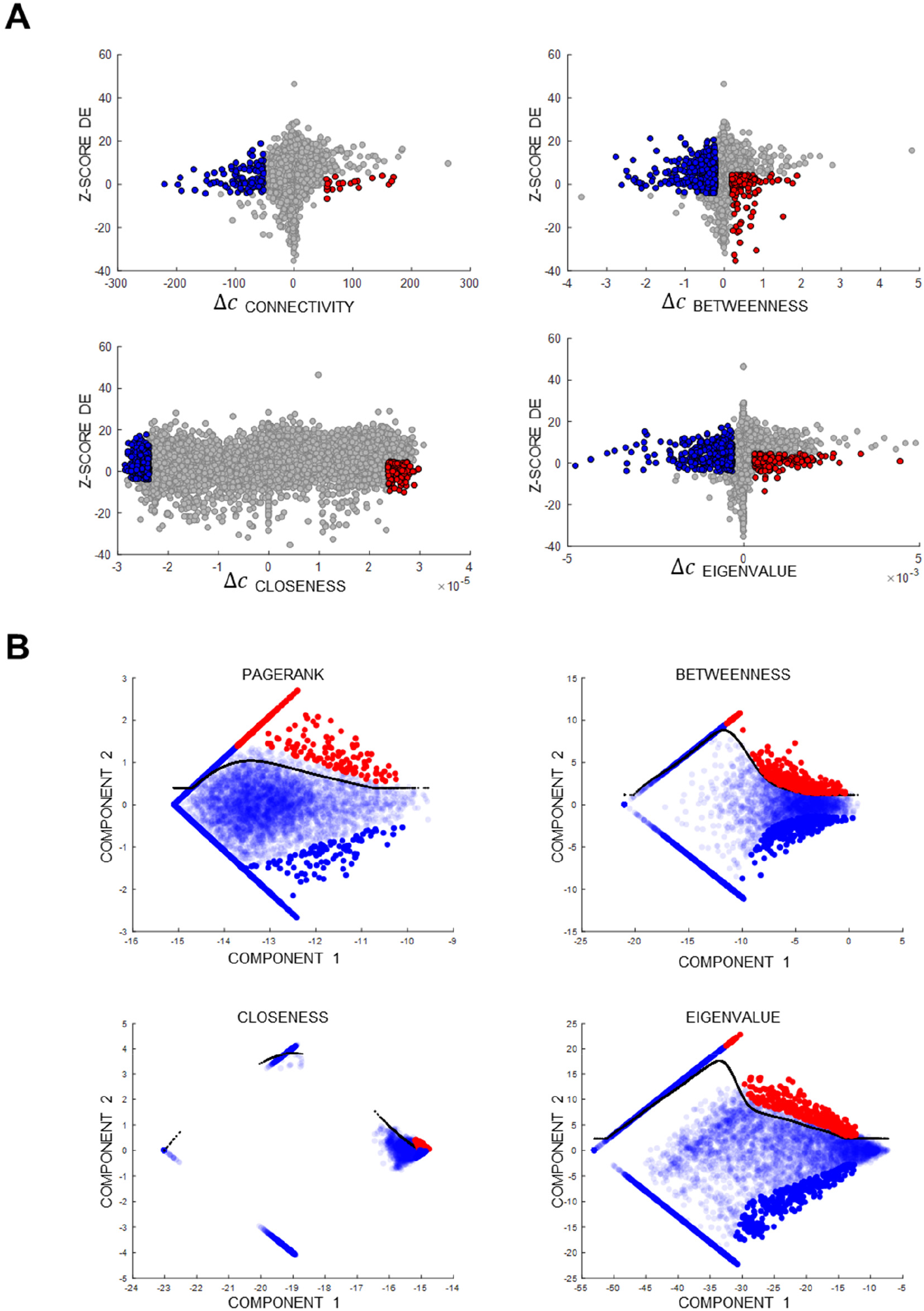
Detection of genes showing changes in centrality. (A) Change in expression as measured by *bigSCale* Z-score (y-axes) compared to the absolute change in the five centralities. (B) Adaptive empirical fitting for the detection of relative changes of centralities.

## References

Aibar, S., González-Blas, C.B., Moerman, T., Huynh-Thu, V.A., Imrichova, H., Hulselmans, G., Rambow, F., Marine, J.-C., Geurts, P., Aerts, J., et al. (2017). SCENIC: single-cell regulatory network inference and clustering. Nature Methods 14, 1083–1086.

Albert, R. (2005). Scale-free networks in cell biology. Journal of Cell Science 118, 4947–4957.

Balaji, S., Iyer, L.M., Aravind, L., and Babu, M.M. (2006). Uncovering a Hidden Distributed Architecture Behind Scale-free Transcriptional Regulatory Networks. Journal of Molecular Biology 360, 204–212.

Chan, T.E., Stumpf, M.P.H., and Babtie, A.C. (2017). Gene Regulatory Network Inference from Single-Cell Data Using Multivariate Information Measures. Cell Syst 5, 251–267.e3.

Chen, S., and Mar, J.C. (2018). Evaluating methods of inferring gene regulatory networks highlights their lack of performance for single cell gene expression data. BMC Bioinformatics 19, 232.

Cheng, H., Jiang, L., Wu, M., and Liu, Q. (2009). Inferring Transcriptional Interactions by the Optimal Integration of ChIP-chip and Knock-out Data. Bioinform Biol Insights 3, 129–140.

Emmert-Streib, F., Dehmer, M., and Haibe-Kains, B. (2014). Gene regulatory networks and their applications: understanding biological and medical problems in terms of networks. Front Cell Dev Biol 2.

Facchetti, G., Iacono, G., and Altafini, C. (2011). Computing global structural balance in large-scale signed social networks. PNAS 108, 20953–20958.

Fiers, M.W.E.J., Minnoye, L., Aibar, S., Bravo González-Blas, C., Kalender Atak, Z., and Aerts, S. (2018). Mapping gene regulatory networks from single-cell omics data. Brief Funct Genomics 17, 246–254.

Ghanevati, M., and Miller, C.A. (2005). Phospho-beta-catenin accumulation in Alzheimer’s disease and in aggresomes attributable to proteasome dysfunction. J. Mol. Neurosci. 25, 79–94.

Guan, M., Keaton, J.M., Dimitrov, L., Hicks, P.J., Xu, J., Palmer, N.D., Wilson, J.G., Freedman, B.I., Bowden, D.W., and Ng, M.C.Y. (2018). An Exome-wide Association Study for Type 2 Diabetes-Attributed End-Stage Kidney Disease in African Americans. Kidney Int Rep 3, 867–878.

Guo, M., Wang, H., Potter, S.S., Whitsett, J.A., and Xu, Y. (2015). SINCERA: A Pipeline for SingleCell RNA-Seq Profiling Analysis. PLoS Comput. Biol. 11, e1004575.

Hamey, F.K., Nestorowa, S., Kinston, S.J., Kent, D.G., Wilson, N.K., and Göttgens, B. (2017). Reconstructing blood stem cell regulatory network models from single-cell molecular profiles. Proc. Natl. Acad. Sci. U.S.A. 114, 5822–5829.

Henriksen, E.J., and Dokken, B.B. (2006). Role of glycogen synthase kinase-3 in insulin resistance and type 2 diabetes. Curr Drug Targets 7, 1435–1441.

Iacono, G., and Altafini, C. (2010). Monotonicity, frustration, and ordered response: an analysis of the energy landscape of perturbed large-scale biological networks. BMC Syst Biol 4, 83.

Iacono, G., Ramezani, F., Soranzo, N., and Altafini, C. (2010). Determining the distance to monotonicity of a biological network: a graph-theoretical approach. IET Systems Biology 4, 223–235.

Iacono, G., Mereu, E., Guillaumet-Adkins, A., Corominas, R., Cuscó, I., RodrÍguez-Esteban, G., Gut, M., Pérez-Jurado, L.A., Gut, I., and Heyn, H. (2018). bigSCale: an analytical framework for big-scale single-cell data. Genome Res. 28, 878–890.

Ishizuka, Y., Nakayama, K., Ogawa, A., Makishima, S., Boonvisut, S., Hirao, A., Iwasaki, Y., Yada, T., Yanagisawa, Y., Miyashita, H., et al. (2014). TRIB1 downregulates hepatic lipogenesis and glycogenesis via multiple molecular interactions. Journal of Molecular Endocrinology 52, 145–158.

Itani, S.I., Ruderman, N.B., Schmieder, F., and Boden, G. (2002). Lipid-induced insulin resistance in human muscle is associated with changes in diacylglycerol, protein kinase C, and IkappaB-alpha. Diabetes 51, 2005–2011.

Keren-Shaul, H., Spinrad, A., Weiner, A., Matcovitch-Natan, O., Dvir-Szternfeld, R., Ulland, T.K., David, E., Baruch, K., Lara-Astaiso, D., Toth, B., et al. (2017). A Unique Microglia Type Associated with Restricting Development of Alzheimer’s Disease. Cell 169, 1276–1290.e17.

Kuo, T., Kim-Muller, J.Y., McGraw, T.E., and Accili, D. (2016). Altered Plasma Profile of Antioxidant Proteins as an Early Correlate of Pancreatic β Cell Dysfunction. J. Biol. Chem. 291, 9648–9656.

Li, F.X., Zhu, J.W., Tessem, J.S., Beilke, J., Varella-Garcia, M., Jensen, J., Hogan, C.J., and DeGregori, J. (2003). The development of diabetes in E2f1/E2f2 mutant mice reveals important roles for bone marrow-derived cells in preventing islet cell loss. Proc. Natl. Acad. Sci. U.S.A. 100, 12935–12940.

Lim, C.Y., Wang, H., Woodhouse, S., Piterman, N., Wernisch, L., Fisher, J., and Göttgens, B. (2016). BTR: training asynchronous Boolean models using single-cell expression data. BMC Bioinformatics 17, 355.

Malecki, M.T., Jhala, U.S., Antonellis, A., Fields, L., Doria, A., Orban, T., Saad, M., Warram, J.H., Montminy, M., and Krolewski, A.S. (1999). Mutations in NEUROD1 are associated with the development of type 2 diabetes mellitus. Nat. Genet. 23, 323–328.

Matsumoto, H., Kiryu, H., Furusawa, C., Ko, M.S.H., Ko, S.B.H., Gouda, N., Hayashi, T., and Nikaido, I. (2017). SCODE: an efficient regulatory network inference algorithm from single-cell RNA-Seq during differentiation. Bioinformatics 33, 2314–2321.

Miller, M.R., Zhang, W., Sibbel, S.P., Langefeld, C.D., Bowden, D.W., Haffner, S.M., Bergman, R.N., Norris, J.M., and Fingerlin, T.E. (2010). Variant in the 3' Region of the kBa Gene Associated With Insulin Resistance in Hispanic Americans: The IRAS Family Study. Obesity (Silver Spring) 18, 555–562.

Moignard, V., Macaulay, I.C., Swiers, G., Buettner, F., Schütte, J., Calero-Nieto, F.J., Kinston, S., Joshi, A., Hannah, R., Theis, F.J., et al. (2013). Characterization of transcriptional networks in blood stem and progenitor cells using high-throughput single-cell gene expression analysis. Nat. Cell Biol. 15, 363–372.

Musso, G., Cassader, M., Bo, S., De Michieli, F., and Gambino, R. (2013). Sterol regulatory element-binding factor 2 (SREBF-2) predicts 7-year NAFLD incidence and severity of liver disease and lipoprotein and glucose dysmetabolism. Diabetes 62, 1109–1120.

Noort, V. van, Snel, B., and Huynen, M.A. (2004). The yeast coexpression network has a small-world, scale-free architecture and can be explained by a simple model. EMBO Reports 5, 280–284.

Omatsu, T., Cepinskas, G., Clarson, C., Patterson, E.K., Alharfi, I.M., Summers, K., Couraud, P.O., Romero, I.A., Weksler, B., Fraser, D.D., et al. (2014). CXCL1/CXCL8 (GROα/IL-8) in human diabetic ketoacidosis plasma facilitates leukocyte recruitment to cerebrovascular endothelium in vitro. Am. J. Physiol. Endocrinol. Metab. 306, E1077–1084.

Papili Gao, N., Ud-Dean, S.M.M., Gandrillon, O., and Gunawan, R. (2017). SINCERITIES: Inferring gene regulatory networks from time-stamped single cell transcriptional expression profiles. Bioinformatics.

Peiris, H., Raghupathi, R., Jessup, C.F., Zanin, M.P., Mohanasundaram, D., Mackenzie, K.D., Chataway, T., Clarke, J.N., Brealey, J., Coates, P.T., et al. (2012). Increased expression of the glucose-responsive gene, RCAN1, causes hypoinsulinemia, β-cell dysfunction, and diabetes. Endocrinology 153, 5212–5221.

Pina, C., Teles, J., Fugazza, C., May, G., Wang, D., Guo, Y., Soneji, S., Brown, J., Edén, P., Ohlsson, M., et al. (2015). Single-Cell Network Analysis Identifies DDIT3 as a Nodal Lineage Regulator in Hematopoiesis. Cell Rep 11, 1503–1510.

Qiu, Y., Zhao, D., Butenschön, V.-M., Bauer, A.T., Schneider, S.W., Skolnik, E.Y., Hammes, H.-P., Wieland, T., and Feng, Y. (2016). Nucleoside diphosphate kinase B deficiency causes a diabetes-like vascular pathology via up-regulation of endothelial angiopoietin-2 in the retina. Acta Diabetol 53, 81–89.

Ravier, M.A., Leduc, M., Richard, J., Linck, N., Varrault, A., Pirot, N., Roussel, M.M., Bockaert, J., Dalle, S., and Bertrand, G. (2014). β-Arrestin2 plays a key role in the modulation of the pancreatic beta cell mass in mice. Diabetologia 57, 532–541.

Sanchez-Castillo, M., Blanco, D., Tienda-Luna, I.M., Carrion, M.C., and Huang, Y. (2018). A Bayesian framework for the inference of gene regulatory networks from time and pseudo-time series data. Bioinformatics 34, 964–970.

Segerstolpe, Å, Palasantza, A., Eliasson, P., Andersson, E.-M., Andréasson, A.-C., Sun, X., Picelli, S., Sabirsh, A., Clausen, M., Bjursell, M.K., et al. (2016). Single-Cell Transcriptome Profiling of Human Pancreatic Islets in Health and Type 2 Diabetes. Cell Metab 24, 593–607.

Shao, W., and Espenshade, P.J. (2012). Expanding roles for SREBP in metabolism. Cell Metab 16, 414–419.

Sonawane, A.R., Platig, J., Fagny, M., Chen, C.-Y., Paulson, J.N., Lopes-Ramos, C.M., DeMeo, D.L., Quackenbush, J., Glass, K., and Kuijjer, M.L. (2017). Understanding Tissue-Specific Gene Regulation. Cell Rep 21, 1077–1088.

Soranzo, N., Ramezani, F., Iacono, G., and Altafini, C. (2012). Decompositions of large-scale biological systems based on dynamical properties. Bioinformatics 28, 76–83.

Tabula Muris Consortium, Overall coordination, Logistical coordination, Organ collection and processing, Library preparation and sequencing, Computational data analysis, Cell type annotation, Writing group, Supplemental text writing group, and Principal investigators (2018). Single-cell transcriptomics of 20 mouse organs creates a Tabula Muris. Nature.

Teran-Garcia, M., Rankinen, T., Rice, T., Leon, A.S., Rao, D.C., Skinner, J.S., and Bouchard, C. (2007). Variations in the four and a half LIM domains 1 gene (FHL1) are associated with fasting insulin and insulin sensitivity responses to regular exercise. Diabetologia 50, 1858–1866.

Thompson, D., Regev, A., and Roy, S. (2015). Comparative analysis of gene regulatory networks: from network reconstruction to evolution. Annu. Rev. Cell Dev. Biol. 31, 399–428.

Tuncay, K., Ensman, L., Sun, J., Haidar, A.A., Stanley, F., Trelinski, M., and Ortoleva, P. (2007). Transcriptional regulatory networks via gene ontology and expression data. In Silico Biol. (Gedrukt) 7, 21–34.

Wanic, K., Krolewski, B., Ju, W., Placha, G., Niewczas, M.A., Walker, W., Warram, J.H., Kretzler, M., and Krolewski, A.S. (2013). Transcriptome analysis of proximal tubular cells (HK-2) exposed to urines of type 1 diabetes patients at risk of early progressive renal function decline. PLoS ONE 8, e57751.

Wei, J., Hu, X., Zou, X., and Tian, T. (2017). Reverse-engineering of gene networks for regulating early blood development from single-cell measurements. BMC Med Genomics 10, 72.

Yang, H., Son, G.W., Park, H.R., Lee, S.E., and Park, Y.S. (2016). Effect of Korean Red Ginseng treatment on the gene expression profile of diabetic rat retina. J Ginseng Res 40, 1–8.

Zappia, L., Phipson, B., and Oshlack, A. (2018). Exploring the single-cell RNA-seq analysis landscape with the scRNA-tools database. PLOS Computational Biology 14, e1006245.

Zeisel, A., Muñoz-Manchado, A.B., Codeluppi, S., Lönnerberg, P., Manno, G.L., Juréus, A., Marques, S., Munguba, H., He, L., Betsholtz, C., et al. (2015). Cell types in the mouse cortex and hippocampus revealed by single-cell RNA-seq. Science 347, 1138–1142.

Zeisel, A., Hochgerner, H., Lönnerberg, P., Johnsson, A., Memic, F., van der Zwan, J., Häring, M., Braun, E., Borm, L.E., La Manno, G., et al. (2018). Molecular Architecture of the Mouse Nervous System. Cell 174, 999–1014.e22.

Zhang, Q., Sun, X., Xiao, X., Zheng, J., Li, M., Yu, M., Ping, F., Wang, Z., Qi, C., Wang, T., et al. (2018). Maternal chromium restriction induces insulin resistance in adult mice offspring through miRNA. Int J Mol Med 41, 1547–1559.

Zhang, Y., Xuan, J., de los Reyes, B.G., Clarke, R., and Ressom, H.W. (2008). Network motif-based identification of transcription factor-target gene relationships by integrating multisource biological data. BMC Bioinformatics 9, 203.

Zhao, J., Xiong, X., Li, Y., Liu, X., Wang, T., Zhang, H., Jiao, Y., Jiang, J., Zhang, H., Tang, Q., et al. (2018). Hepatic F-Box Protein FBXW7 Maintains Glucose Homeostasis Through Degradation of Fetuin-A. Diabetes 67, 818–830.

Zhu, Z., Tong, X., Zhu, Z., Liang, M., Cui, W., Su, K., Li, M.D., and Zhu, J. (2013). Development of GMDR-GPU for gene-gene interaction analysis and its application to WTCCC GWAS data for type 2 diabetes. PLoS ONE 8, e61943.

